# Targeted complement inhibition at synapses prevents microglial synaptic engulfment and synapse loss in demyelinating disease

**DOI:** 10.1101/841601

**Authors:** Sebastian Werneburg, Jonathan Jung, Rejani B. Kunjamma, Seung-Kwon Ha, Nicholas J. Luciano, Cory M. Willis, Guangping Gao, Stephen J. Crocker, Brian Popko, Daniel S. Reich, Dorothy P. Schafer

**Affiliations:** Department of Neurobiology, Brudnik Neuropsychiatric Research Institute, University of Massachusetts Medical School, Worcester, MA 01605, USA; Department of Neurology, University of Chicago, Chicago, Illinois 60637, USA; Translational Neuroradiology Section, National Institute of Neurological Disorders and Stroke, National Institutes of Health, Bethesda, MD 20892, USA; Department of Neuroscience, University of Connecticut School of Medicine, Farmington, CT 06032, USA; Horae Gene Therapy Center, University of Massachusetts Medical School, Worcester, MA 01605, USA; Li Weibo Institute for Rare Diseases Research, University of Massachusetts Medical School, Worcester, MA 01605, USA; Department of Microbiologic and Physiological Systems, University of Massachusetts Medical School, Worcester, MA 01605, USA

**Author notes:** Lead Contact/Correspondence.

**Keywords:** multiple sclerosis, neuroinflammation, microglia, synapse, neurodegeneration, demyelination, complement, gene therapy

## Abstract

Multiple sclerosis (MS) is a demyelinating, autoimmune disease of the central nervous system. While work has focused on axon loss in MS, far less is known about synaptic changes. Here, in striking similarity to other neurodegenerative diseases, we identify in postmortem human MS tissue and in nonhuman primate and mouse MS models profound synapse loss and microglial synaptic engulfment. These events can occur independently of local demyelination, neuronal degeneration, and peripheral immune cell infiltration, but coincide with gliosis and increased localization of complement component C3, but not C1q, at synapses. Finally, we use AAV9 to overexpress the complement inhibitor Crry at activated C3-bound synapses in mice and demonstrate robust protection of synapses and visual function. These results mechanistically dissect synapse loss as an early pathology in MS. We further provide a novel gene therapy approach to prevent synapse loss by microglia, which may be broadly applicable to other neurodegenerative diseases.

## Introduction

Multiple sclerosis (MS) is a neurological disease of the central nervous system (CNS) affecting more than 2 million people worldwide (2019). The disease is typically characterized by recurrent episodes of inflammatory demyelination with a relapsing-remitting course, which can be accompanied by neurodegeneration (Reich et al., 2018, Amato et al., 2010). A subset of patients initially present with or develop a progressive, chronic neurodegenerative disease with significant synapse loss and CNS atrophy, termed progressive MS (Mahad et al., 2015). Current FDA and EMA-approved disease-modifying therapies, which target inhibition of peripheral immune attack on the CNS in MS, are increasingly effective at reducing episodes of inflammatory demyelination and neurological disability (Mahad et al., 2015, Weideman et al., 2017). However, the neurodegenerative process, particularly for patients with progressive MS, has proven significantly more challenging to decelerate (Ciotti and Cross, 2018). Similar to other neurodegenerative diseases, there is no clear mechanistic understanding of why some patients develop profound degeneration and disability. Therefore, studying neurodegeneration in MS may offer a unique opportunity to capture early phases of the degenerative process, which may be broadly applicable to other CNS diseases and could lead to novel therapeutic strategies to meet an urgent clinical need.

Synapse loss has recently emerged as an early and likely key feature underlying circuit dysfunction in many neurodegenerative diseases, including Alzheimer’s disease and other dementias (Selkoe et al., 2008, Mucke and Selkoe, 2012, Selkoe, 2002, Milnerwood and Raymond, 2010, Yoshiyama et al., 2007, Coleman et al., 2004, Forner et al., 2017, Tyebji and Hannan, 2017, Henstridge et al., 2016). However, compared to other diseases, far less is known regarding how synaptic connections are affected in MS. The vast majority of research aimed at treating neurodegenerative aspects of MS have focused on mechanisms of de- and remyelination, as well as axon de- and regeneration (Lassmann, 2018, Lassmann, 2010, Mahad et al., 2015, Dutta and Trapp, 2011, Reich et al., 2018). From the few studies assessing synaptic changes in postmortem MS tissue, synapse loss has been observed in the hippocampus and normal appearing gray matter in the cortex (Dutta et al., 2011, Jurgens et al., 2016). Similar results have been found in rodent models of demyelinating disease. For example, in a cuprizone model of demyelination, a significant decrease in excitatory synapses was observed in the visual thalamus concomitant with reactive gliosis and subcortical demyelination (Araujo et al., 2017). In another study, experimental autoimmune encephalomyelitis (EAE)-induced demyelination in mice resulted in a ∼28% decrease in PSD-95-positive postsynaptic densities in the hippocampus, which was observed in the absence of hippocampal demyelination, but in the presence of reactive, phagocytic microglia (Bellizzi et al., 2016). A common feature in all these human and animal model studies is reactive gliosis, including pronounced increases in inflammatory microglia, a resident CNS macrophage (Lassmann, 2018, Lassmann, 2010, Mahad et al., 2015, Voet et al., 2018). However, it remains unclear if and how these inflammatory glial cells could modulate synaptic connectivity in demyelinating disease.

Microglia have recently been identified as key regulators of synaptic connectivity in the healthy and diseased brain. During development, microglia regulate synaptic pruning by engulfing and removing a subset of synapses that initially form in excess. One key mechanism is classical complement cascade-dependent phagocytic signaling (Stevens et al., 2007, Schafer et al., 2012). In the peripheral immune system, components of the classical complement cascade, C1q and C3, bind the surface of invading pathogens, cellular debris, etc., leading to clearance by professional phagocytes that express complement receptors. Similarly, in the developing rodent visual thalamus, C1q and C3 localize to synapses (Stevens et al., 2007). Microglia expressing complement receptor 3 (CR3), a C3 receptor, then engulf these complement-associated synapses. Mice deficient in either CR3 expressed by microglia, C3, or C1q have a ∼50% decrease in their ability to engulf and remove synapses (Schafer et al., 2012, Bialas and Stevens, 2013). Interestingly, this classical complement cascade-mediated phagocytic signaling has now been identified to be aberrantly upregulated in mouse models of Alzheimer’s disease, frontotemporal dementia, and West Nile Virus infection, leading to synapse loss (Hong et al., 2016, Vasek et al., 2016, Lui et al., 2016). In MS patients, complement proteins are elevated systemically and in the CNS (Aeinehband et al., 2015, Ingram et al., 2012, Ingram et al., 2009, Watkins et al., 2016), and recent work has suggested that C1q and C3 colocalize with synaptic proteins in postmortem MS brains (Michailidou et al., 2015). However, it remains elusive if complement and/or microglia are necessary for synaptic changes in MS.

In the current study, we use the retinogeniculate system to study synaptic changes in demyelinating disease. The retinogeniculate system is comprised of retinal ganglion cells (RGCs), which are neurons in the retina that extend their axons via the optic nerve and tract and synapse onto relay neurons within the lateral geniculate nucleus (LGN) of the thalamus. This circuit was chosen because even subtle synaptic changes are easily detected by immunohistochemical methods (Schafer et al., 2016, Schafer et al., 2012, Hong et al., 2014). In addition, because the anterior visual pathway is commonly affected in MS, >50% of patients experience inflammation of the optic nerve (i.e. optic neuritis) and visual dysfunction (Toosy et al., 2014, Reich et al., 2018). Using postmortem human MS tissue, a preclinical nonhuman primate model of MS, and two different rodent models of demyelinating disease, we investigated early synapse changes in the retinogeniculate system. We first show profound synapse loss in the LGN from human MS tissue and in all animal MS models and identify that microglia, but not astrocytes, are engulfing synapses across species. Using rodent models, we further show that these synaptic changes can occur independent of neuronal cell death, axon degeneration, significant demyelination, or peripheral immune cell infiltration. To identify a molecular mechanism underlying this synapse loss, we focused on the complement cascade and provide evidence for activation of the alternative, but not the classical, complement cascade at synapses in demyelinating disease. We then use an adeno-associated viral (AAV) approach to overexpress the C3 inhibitor Crry at sites of synaptic C3 activation in the retinogeniculate circuit and demonstrate robust protection from synapse engulfment by microglia and synapse loss. The rescue of structural synapses with this AAV approach was specific to the retinogeniculate circuit and restored function as assessed by visual acuity. Together, our data provide evidence across multiple species that local inflammatory microglia and the alternative complement cascade mediate synapse loss and functional decline in demyelinating disease. With our data demonstrating that targeted inhibition of activated C3 protects synapses, we further uncover a new therapeutic strategy to prevent synapse loss at specific synapses and preserve visual function in neurodegeneration.

## Results

### Profound synapse loss and microglial engulfment of presynaptic terminals in MS patients and in a preclinical marmoset EAE model

To first interrogate synaptic changes that are clinically relevant to MS, we assessed synaptic connectivity in postmortem LGN from MS patients and control patients without neurological disease (Table S1). Using anti-vesicular glutamate transporter 2 (VGluT2) immunostaining, a marker specific to retinogeniculate presynaptic inputs, and confocal imaging, we observed a significant decrease in retinogeniculate presynaptic terminals within the LGN of MS patients compared to controls (Fig. 1A). Co-labeling with the microglia/macrophage marker Iba1 and 3D-surface rendering further revealed that significant amounts of VGluT2^+^-terminals were engulfed within Iba1^+^-cells (Fig. 1B). These results are consistent with previous work showing significant degeneration of the thalamus and deep gray matter early in the disease course of MS, which accurately predicted subsequent disease severity (Eshaghi et al., 2018, Zivadinov et al., 2013). This is also in line with studies showing decreased presynaptic inputs in the hippocampus and cortex of postmortem MS tissue (Dutta et al., 2011, Jurgens et al., 2016) and studies demonstrating that microglia engulf and eliminate synaptic components in other models of neurodegenerative disease (Hong et al., 2016, Paolicelli et al., 2017, Lui et al., 2016, Vasek et al., 2016).

**Figure 1:**
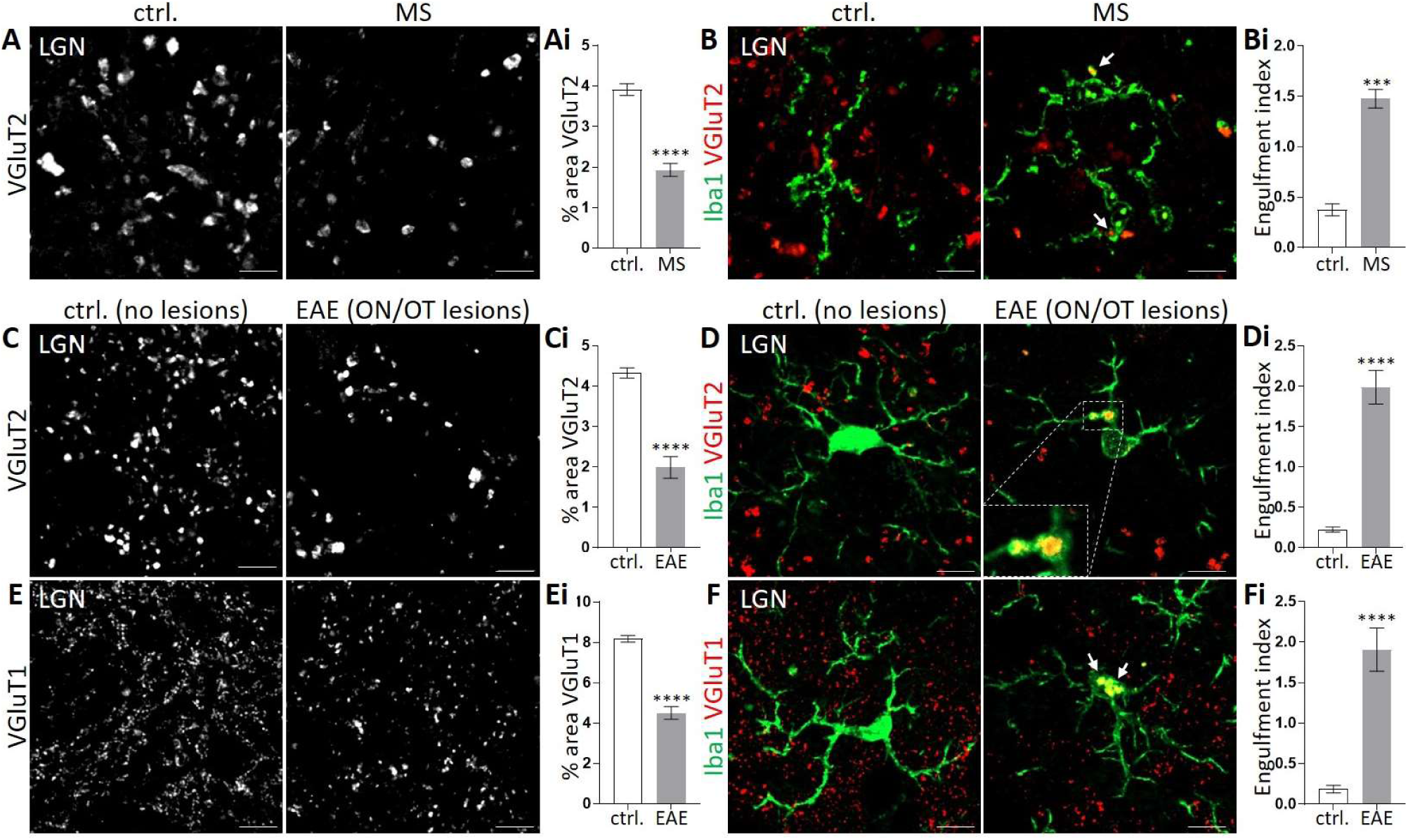
Synapse loss and engulfment of presynaptic terminals in the LGN of MS patients and a preclinical marmoset EAE model. (**A,B**) Coronal sections of the LGN from human controls (ctrl.) without neurological disease and from MS cases demonstrate a decrease in the density of (**Ai**) VGluT2^+^-retinogeniculate presynaptic inputs (red) and (**Bi**) an increase in engulfment (arrows) by Iba1-labeled microglia/macrophages (green) in MS tissue. (**Ai**) n=5 for each condition, (**Bi**) n=3 ctrl./5 MS. Scale bars, 10 μm. (**C-F**) Representative images of the LGN from non-EAE or EAE animals with no detectable lesions (control, no lesions) and marmosets that developed EAE with demyelinating lesions in the optic nerve and tract (EAE with ON/OT lesions). (**Ci**) Similar to human tissue, there is a significant decrease in the density of VGluT2^+^-inputs in the LGN from marmosets that developed ON/OT lesions following EAE compared to controls. (**Di**) Significant amounts of VGluT2^+^-presynaptic RGC inputs were also detected within Iba1-labeled microglia/macrophages in the LGN of marmosets that developed ON/OT lesions following EAE compared to controls. (**D**) Inset shows phagocytic cups within Iba1^+^-cell (green), which contain VGluT2^+^-retinogeniculate synaptic material (red). No phagocytic cups were detected in controls. (**E,F**) VGluT1 staining reveals reduced density of (**E**) VGluT1^+^-corticothalamic presynaptic inputs and (**F**) increased engulfment of VGluT1^+^-synaptic material within Iba1^+^-cells in the LGN of marmosets that developed ON/OT lesions following EAE. (**F**) Arrows denote engulfed VGluT1^+^-corticothalamic inputs. (**C-F**) n=6. Scale bars, 10 μm. Data represent mean ± SEM, significant differences with ***P < 0.001, ****P < 0.0001, t-test. See also Fig. S1 & Table S1+2.

We next sought to corroborate these findings in a preclinical nonhuman primate model of MS, experimental autoimmune encephalomyelitis (EAE) in common marmosets induced by immunization with human white matter homogenate (Lee et al., 2018). This model is particularly powerful as it has many of the pathological features present in human MS (Absinta et al., 2016). Similar to MS patient LGN, we found a ∼2-fold reduction in the density of retinogeniculate terminals and an increased localization of VGluT2 within microglia (Fig. 1C,D) in the LGN from animals that developed EAE with demyelinating lesions in the optic nerve and tract by MRI (Table S2). This is in contrast to either EAE or non-EAE animals with no detectable lesions in the optic nerve and tract (Table S2), which were used as controls. Many of the engulfed VGluT2^+^-retinogeniculate terminals were detected within microglial phagocytic cups, which were not present in microglia from control animals (Fig. 1D). A recent MRI study indicated that damage of the LGN in MS patients results from anterograde degeneration from the retina and retrograde degeneration from the visual cortex (Papadopoulou et al., 2019). To determine if loss of presynaptic inputs and engulfment by microglia is specific to retinogeniculate terminals or if synaptic connectivity within the LGN is affected more broadly, we assessed the density of corticothalamic inputs by immunostaining against VGluT1 in marmosets. Similar to retinogeniculate terminals, we also found a significant decrease in the density of VGluT1^+^-corticothalamic inputs in the LGN and detected significant amounts of VGluT1 within microglia in marmosets with lesions in the optic nerve and tract following EAE (Fig. 1E,F). These data provide the first evidence that reactive microglia/macrophages engulf and eliminate presynaptic inputs within the retinogeniculate circuit in demyelinating disease

### Synapse loss can occur independent of significant demyelination, cell death, or axon degeneration

In the human MS cases, patients have had protracted disease with significant bouts of demyelination. In the marmoset EAE model, the animals also showed signs of demyelination in the LGN and optic tract by MRI (Table S2), which we confirmed in postmortem tissue (Fig. S1A,B). To determine if synapse loss occurred independent of demyelination and to gain a deeper understanding of the cellular and molecular mechanisms underlying synapse loss in demyelinating disease, we next interrogated retinogeniculate synaptic changes in the MOG_35-55_-induced mouse EAE model prior to peak clinical score compared to Complete Freund’s Adjuvant (CFA)-treated controls (Crocker et al., 2006). Mice analyzed here displayed moderate clinical symptoms (average scores: 1.35±0.35) that were typically observed between day 10-12 post-immunization (average: 10.8±0.4) (Fig. S2A). Similar to human and marmoset data (Fig. 1), we detected a significant, >50% reduction in VGluT2^+^- and VGluT1^+^-presynaptic inputs within the LGN at this earlier stage of mouse EAE (Fig. 2A,B). This was accompanied by a pronounced reduction in the density of structural synapses (co-localized presynaptic VGluT2 or VGluT1 with postsynaptic PSD-95 or Homer1). However, this synapse loss in EAE was mainly attributed to a presynaptic effect, as the densities of postsynaptic PSD-95 and Homer1 were relatively unaltered (Fig. 2A,B).

**Figure 2:**
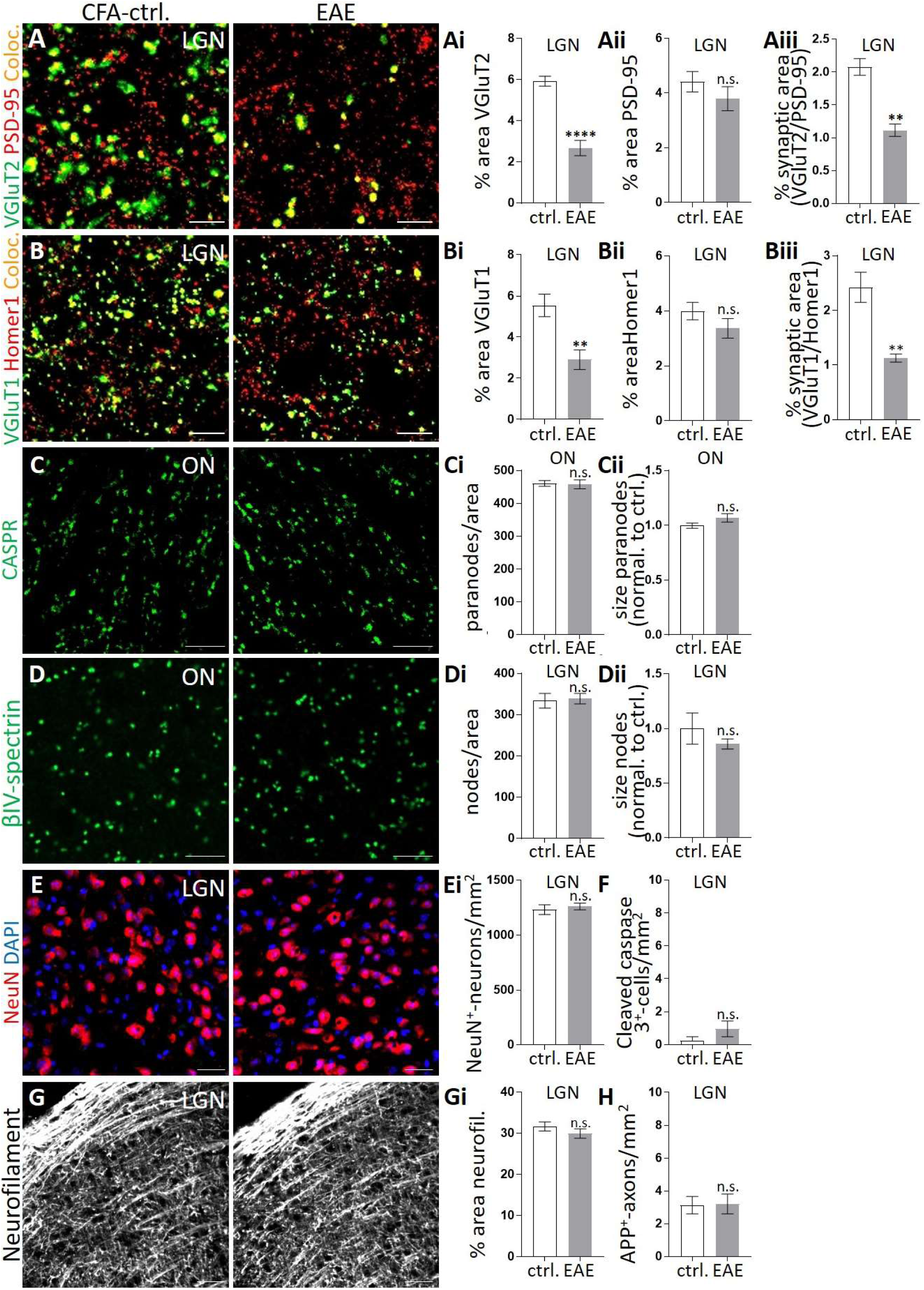
Synapse loss in the LGN can occur in the absence of significant demyelination, cell death, or axon degeneration in early stages of mouse EAE. (**A,B**) Representative images of the LGN from CFA-control and EAE-induced mice immunostained against (**A**) presynaptic VGluT2 (green) and postsynaptic PSD-95 (red) or (**B**) presynaptic VGluT1 (green) and postsynaptic Homer1 (red). Quantitative analysis shows markedly reduced densities of presynaptic (**Ai**) VGluT2 and (**Bi**) VGluT1, and unaltered densities of postsynaptic (**Aii**) PSD-95 and (**Bii**) Homer1 in mouse EAE LGN compared to CFA-injected controls. This was also reflected in a loss of structural (**Aiii**) retinogeniculate and (**Biii**) corticothalamic synaptic areas (defined as colocalized pre and postsynaptic markers (Coloc.) in EAE LGN compared to controls. Scale bars, 5 μm. (**C,D**) Analysis of the optic nerve (ON) of the same CFA-control and EAE-treated mice. Representative images of (**C**) anti-CASPR-labeled paranodes and (**D**) anti-βIV-spectrin-labeled nodes of Ranvier. Quantification demonstrates no changes in the (**Ci,Di**) density and no alterations in (**Cii,Dii**) size (normalized to controls) of (**Ci,Cii**) paranodes and (**Di,Dii**) nodes of Ranvier following EAE. Scale bars, 10 μm. (**E**) Representative images of immunostaining in the LGN for the neuronal maker NeuN (red), along with a DAPI counterstain to label all nuclei (blue). Quantification demonstrates no differences in the density of (**Ei**) NeuN^+^-neurons or (**F**) cleaved caspase 3^+^-cells, a marker of cell death, between groups. (**G**) Representative images of neurofilament (neurofil.)-labeled axons. Quantification demonstrates no changes in the density of (**Gi**) neurofilament^+^-axons and (**H**) no induction of amyloid precursor protein (APP), a marker of axonal degeneration in EAE tissue compared to controls. Scale bars, 20 μm. (**A-H**) n=4. Data represent mean ± SEM, significant differences with **P < 0.01, ****P < 0.0001, t-test. See also Fig. S2-4.

We next addressed whether synapse loss was a secondary effect to other brain pathology or whether it occurred independent of demyelination, cell death, and/or axon degeneration. Consistent with the latter, while a significant increase in peripheral CD3^+^-T-cells, CD45^+^-leukocytes (Fig. S2B,C), reactive microglia, and GFAP^+^-astrocytes (Fig. S2D,E) were observed, we detected no significant changes in the myelin sheath (Figs. 2C,D; S3). This is consistent with capturing events at early stages of EAE where the animals had moderate clinical scores (average scores: 1.35±0.35) at the time of sacrificing. This lack of demyelination in the retinogeniculate system was assessed by measuring for disruptions in paranodal junctions and nodes of Ranvier, which have previously been described as early events in demyelination (Wolswijk and Balesar, 2003, Howell et al., 2006). No significant alterations in the density or size of CASPR^+^-paranodal junctions or βIV-spectrin^+^-nodes of Ranvier were observed in the retinogeniculate circuit at this stage of EAE (Fig. 2C,D). Further supporting that myelin sheaths are unaltered at the time point when we observe synapse loss, no detectable changes in myelin proteins (MOG, MBP, and MAG) were observed in the optic tract, the LGN, or in longitudinal sections along the length of the optic nerve in EAE compared to CFA-controls (Fig. S3). We next assessed cell death and axon degeneration. Similar to no significant changes in the myelin sheath, there were no significant changes in the density of NeuN^+^-neurons or neurofilament^+^-axons and no significant increases in the number of cleaved caspase 3^+^-apoptotic cells or amyloid precursor protein (APP) accumulations in axons within the retinogeniculate circuit at these early phases in the mouse EAE model (Figs. 2E-H; S4). We also found no significant alterations in neurons and axons within the LGN of the marmoset EAE model upon further examination (Fig. S1C-E). Together, these data are consistent with synapse loss occurring concomitant with local inflammation (reactive glia and peripheral immune cell infiltrates), but that it can occur independent of demyelination, cell death, and axon degeneration (Hong et al., 2016, Paolicelli et al., 2017, Wishart et al., 2006, Yoshiyama et al., 2007).

### Microglia, but not astrocytes, engulf synapses in animal models of MS

To further examine whether local inflammatory gliosis was a contributing factor to early synapse loss, we tested whether reactive microglia engulfed presynaptic terminals in the mouse EAE model. We labeled microglia with anti-P2RY12, which distinguishes resident microglia from peripheral infiltrating macrophages (Butovsky et al., 2014, Jordao et al., 2019), and microglial lysosomes with anti-CD68 to assess engulfed presynaptic terminals within CD68^+^-microglial lysosomes using high-resolution confocal imaging and 3D-surface rendering. Similar to data from human and marmoset (Fig. 1), we found a ∼10-fold increase in the engulfment of VGluT2^+^- and VGluT1^+^-presynaptic inputs within microglial lysosomes in the LGN in EAE mice (Fig. 3A,B). In contrast, only minimal amounts of presynaptic inputs were detected within microglial lysosomes in CFA-control animals. We found no evidence for engulfment of postsynaptic compartments in EAE mice (Fig. 3C). This finding is consistent with other reports showing that microglia predominantly engulf presynaptic inputs (Schafer et al., 2012, Schafer et al., 2014, Weinhard et al., 2018), and our findings that the density of postsynaptic compartments is relatively unaltered in EAE mice (Fig. 2A,B). In addition to microglia, astrocytes have been shown to engulf synaptic material during postnatal development of the visual system (Chung et al., 2013). To test if reactive astrocytes similarly engulf synaptic material following EAE, we stained astrocytes in the LGN with an antibody against ALDH1L1 and assessed engulfment of VGluT2 and VGluT1 into LAMP2^+^-lysosomes. Surprisingly, no engulfment of VGluT2 or VGluT1 was detected within astrocytes in mouse or marmoset EAE models (Fig. 3D-F). These data suggest that while astrocytes engulf synapses during developmental synaptic pruning, reactive astrocytes are not contributing to the engulfment of presynaptic terminals in demyelinating disease. Our data are most consistent with microglia as primary mediators of synapse elimination in this demyelinating disease context.

**Figure 3:**
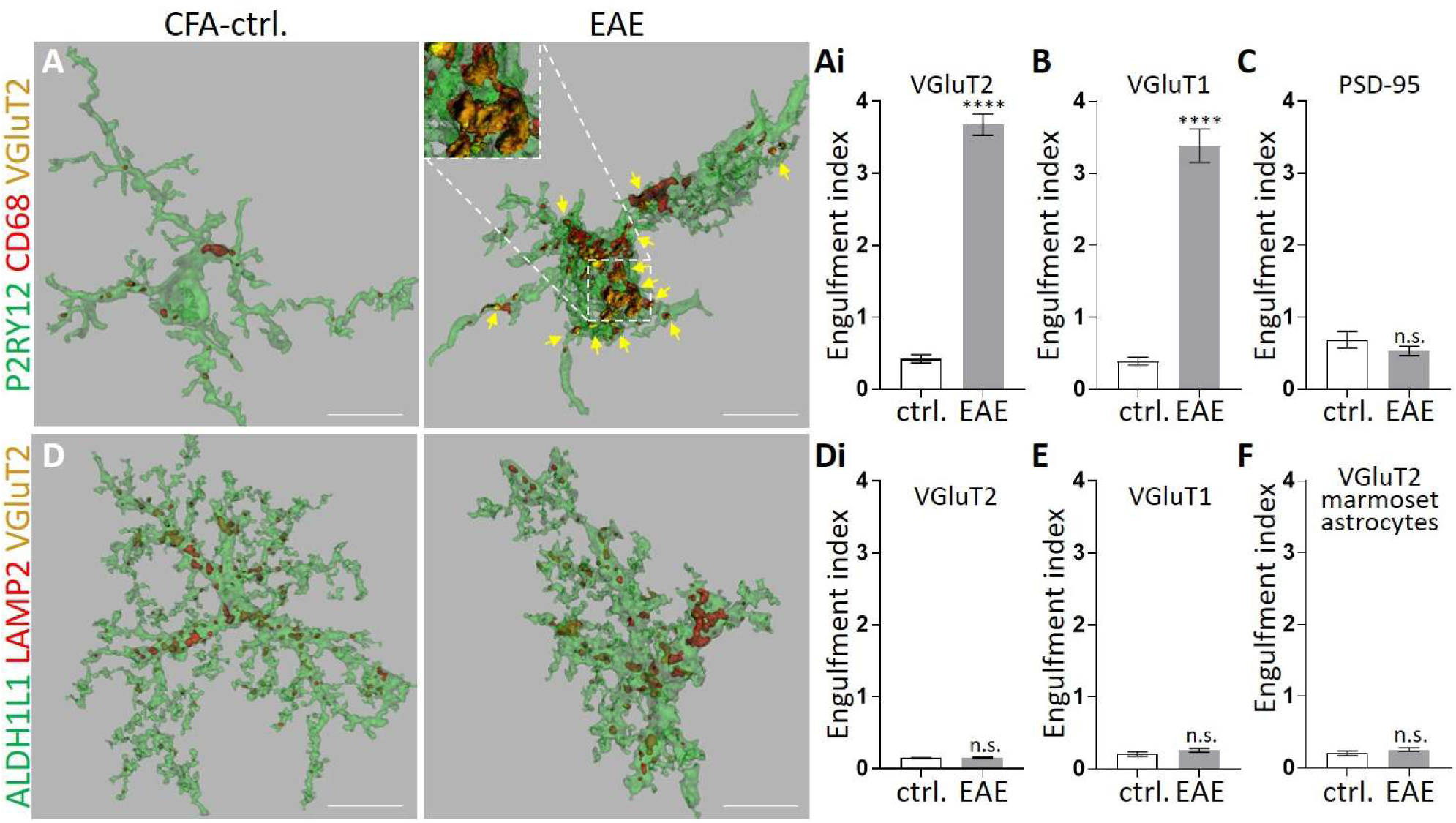
Microglia, but not astrocytes, engulf presynaptic inputs at early stages of mouse EAE. (**A-F**) Synapse engulfment was analyzed in the LGN from the same mouse EAE tissue in which we observed synapse loss independent from demyelination, axon degeneration, and cell death. (**A,B**) Representative 3D-surface rendering of P2RY12^+^-microglia (green) reveals VGluT2^+^-retinogeniculate inputs (yellow, arrows, insert) within CD68-labeled microglial lysosomes (red) in EAE mice (right) compared to CFA-controls (left). Microglia engulf significantly more VGluT2^+^-retinogeniculate (**Ai**) and VGluT1^+^-corticothalamic presynaptic inputs (**B**) in EAE mice compared to controls. (**C**) There was no evidence for engulfment of postsynaptic density protein-95 (PSD-95)-labeled postsynaptic compartments. (**D**) Representative 3D-surface rendering of ALDH1L1^+^-astrocytes (green) reveals little to no detection of VGluT2^+^-retinogeniculate inputs (yellow) within LAMP2-labeled lysosomes (red) in the LGN of EAE or control mice. (**Di,E**) Quantification shows no significant increase in (**Di**) VGluT2 or (**E**) VGluT1 within reactive astrocytes in EAE mice. (**F**) Similar to mouse EAE, ALDH1L1-labeled astrocytes show no significant increase in engulfed VGluT2^+^-retinogeniculate inputs within LAMP2^+^-lysosomes in the marmoset EAE model. (**A-E**) n=4. (**F**) n=6. (**A,D**). Scale bars, 10 μm. Data represent mean ± SEM, significant differences with ****P < 0.0001, t-test.

### Microglial synaptic engulfment and synapse loss can occur independent of peripheral immune cell infiltration

While we observed no detectable induction of demyelination, cell death, or axon degeneration in the mouse EAE model, a significant increase in infiltrating peripheral T-cells and leukocytes was observed. To investigate the influence of peripheral immune cells on microglia-mediated synapse engulfment and synapse loss, we assessed *Plp1-CreER*^*T*^;*ROSA26-EGFP-DTA* (diphtheria toxin A (DTA)) mice compared to ROSA26-EGFP-DTA control mice (ctrl.). In this paradigm, 5-7-week-old DTA and ctrl. mice receive tamoxifen injections. In DTA mice, this induces the expression of diphtheria toxin A in mature oligodendrocytes, which results in demyelination within several weeks post-injection (Fig. 4A) (Traka et al., 2010, Traka et al., 2016). While this model is not conducive to dissecting whether synapse loss is independent of demyelination, there is very minimal peripheral immune cell infiltration, allowing us to test whether synapse loss could occur independent from peripheral immune cell involvement. We first confirmed progressive demyelination in the retinogeniculate circuit as indicated by a reduction of MOG, MBP, and MAG in the visual system (Fig. S5A-F). There was no significant infiltration of CD3^+^-T-cells and CD45^+^-leukocytes at 21dpi. By 35 dpi, there was a subtle increase in peripheral immune cell populations compared to controls (Fig. 4C,D); however, the number of infiltrates was >10-fold less than numbers observed in EAE mice (Fig. S2B,C). In the absence of large amounts of peripheral immune infiltrates, we did observe significant microgliosis, exemplified by a progressive increase in the number of microglia that had decreased expression of P2RY12, which also acquired a more reactive, amoeboid-like morphology over the course of demyelination (Figs. 4B; S5G). We then measured synapse engulfment by reactive microglia. As observed in EAE models, we detected significant amounts of engulfed VGluT2^+^- and VGluT1^+^-presynaptic terminals within microglia in the DTA mice compared to controls (Fig. 4E,F), while no synaptic material was detected within astrocytes (Fig. 4G). Notably, microglial engulfment of presynaptic inputs could be detected at earlier stages of demyelination (21dpi, Fig. 4E,F) when no significant recruitment of peripheral immune cells to the LGN was detected (Fig. 4C,D) and prior to significant synapse loss at 35 dpi (Fig. 4H,I). In agreement with the EAE models, engulfment and elimination of presynaptic terminals occurred in the absence of cell death, axonal degeneration, or decreases in postsynaptic compartments in DTA mice (Figs. 4H,I; S5H-L). These results further support that microglia engulf and eliminate synapses in demyelinating disease and suggest that this biology can occur independent of peripheral immune cell involvement.

**Figure 4:**
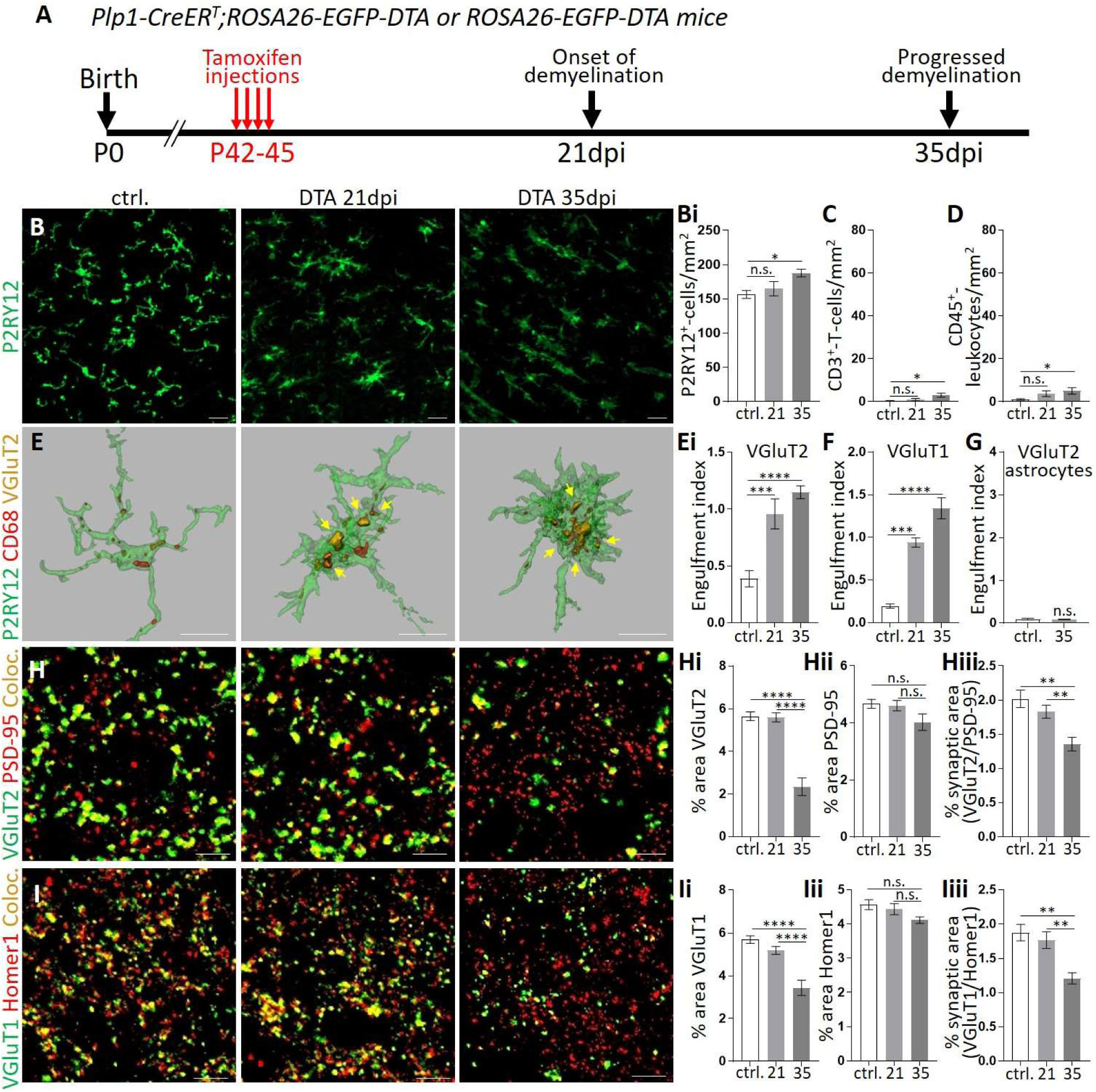
Microglial synaptic engulfment and synapse loss in the DTA model of demyelination occurs with minimal infiltration of peripheral immune cells. (**A**) Timeline for tamoxifen-induced genetic depletion of mature oligodendrocytes in *Plp1-CreER*^*T*^;*ROSA26-EGFP-DTA* (DTA) mice. Tamoxifen-injected ROSA26-EGFP-DTA mice were used as controls (ctrl.). The LGN of mice was analyzed at 21 (onset of demyelination) and 35 (progressed demyelination) days post induction (21dpi and 35dpi, respectively). (**B**) Immunostaining against P2RY12 reveals progressive microgliosis, exemplified by reduced levels of P2RY12 (see also Fig. S5G), progressive acquisition of a reactive, amoeboid-like morphology, and increasing numbers of (**Bi**) P2RY12^+^-microglia. (**C,D**) Quantification of CD3^+^-T-cells and CD45^+^-leukocytes in the LGN shows minimal infiltration of peripheral immune cells. (**E**) Representative 3D-surface rendering of P2RY12^+^-microglia (green) reveals engulfed VGluT2^+^-retinogeniculate inputs (yellow, arrows) within CD68-labeled microglial lysosomes (red) in DTA mice. Scale bars, 10 μm. (**Ei,F**) Microglia engulf significantly more (**Ei**) VGluT2^+^-retinogeniculate and (**F**) VGluT1^+^-corticothalamic presynaptic inputs within CD68^+^-lysosomes in DTA mice compared to controls. (**G**) No engulfment of VGluT2^+^-retinogeniculate inputs was detected within LAMP2-labeled lysosomes of ALDH1L1^+^-astrocytes in DTA mice. (**H,I**) Representative confocal images of the LGN from control and DTA mice immunostained against (**H**) presynaptic VGluT2 (green) and postsynaptic PSD-95 (red) or (**I**) presynaptic VGluT1 (green) and postsynaptic Homer1 (red). There was a significant decrease in presynaptic (**Hi**) VGluT2 and (**Ii**) VGluT1 input density in the LGN of DTA mice by 35dpi. (**Hii,Iii**) In contrast, postsynaptic markers were relatively spared in DTA mice. (**Hiii,Iiii**) Consistent with reduced presynaptic terminal markers, there was a significant reduction in structural synaptic area in the LGN of DTA mice. Structural synapses were defined as colocalized (Coloc., yellow) pre- and postsynaptic markers. (**B-F**) n=9 ctrl./5 DTA-21dpi/6 DTA-35dpi. (**G**) n=4. (**H,I**) n=5 ctrl./4 DTA. Scale bars, (**B**) 20 μm (**H,I**) 5 μm. Data represent mean ± SEM, significant differences with *P < 0.05, **P < 0.01, ***P < 0.001, ****P < 0.0001, ANOVA with Tukey’s *post hoc* test. See also Fig. S5.

### Complement component C3, but not C1q, localizes to synapses in animal models of MS

Previous work in mice has shown that microglial engulfment and elimination of synapses in development and in models of neurodegeneration is mediated by the classical complement cascade. In this cascade, the initiating molecule C1q and downstream C3 localize to synapses and microglia that express the receptor for C3 (CR3) engulf and eliminate synapses (Schafer et al., 2012, Hong et al., 2016, Stevens et al., 2007). Interestingly, these molecules have been shown to increase in other CNS regions in EAE and MS patient tissue (Nataf et al., 2000), but it is unknown if complement mediates synapse loss in these contexts. We first assessed levels of C1q and C3 in the retinogeniculate system in mouse EAE. This analysis revealed a significant increase in both complement factors following EAE (Fig. 5A-D). However, despite increases in C1q throughout the LGN, it did not colocalize with retinogeniculate or corticothalamic presynaptic terminals in EAE (Fig. 5A,B). In contrast, complement factor C3 showed striking colocalization with both presynaptic terminal markers and enrichment at synaptic compartments (Fig. 5C,D). Further, the same observations were made in the marmoset EAE and mouse DTA models (Figs. 5E,F; S6). In the DTA model, C3 colocalization with presynaptic markers could be detected even at earlier stages of disease (21dpi), when microglia engulf synaptic terminals but prior to significant synapse loss (Fig. S6). These data support a complement-dependent model by which microglia engulf and eliminate synapses (Hong et al., 2016, Paolicelli et al., 2017, Lui et al., 2016, Vasek et al., 2016). However, unlike developmental and other neurodegenerative disease contexts, synapse loss in demyelinating disease may be independent of C1q localization to synapses. Instead, our data support the involvement of the alternative complement cascade, which bypasses C1q and works directly through synaptically localized, activated C3 to mediate synapse loss.

**Figure 5:**
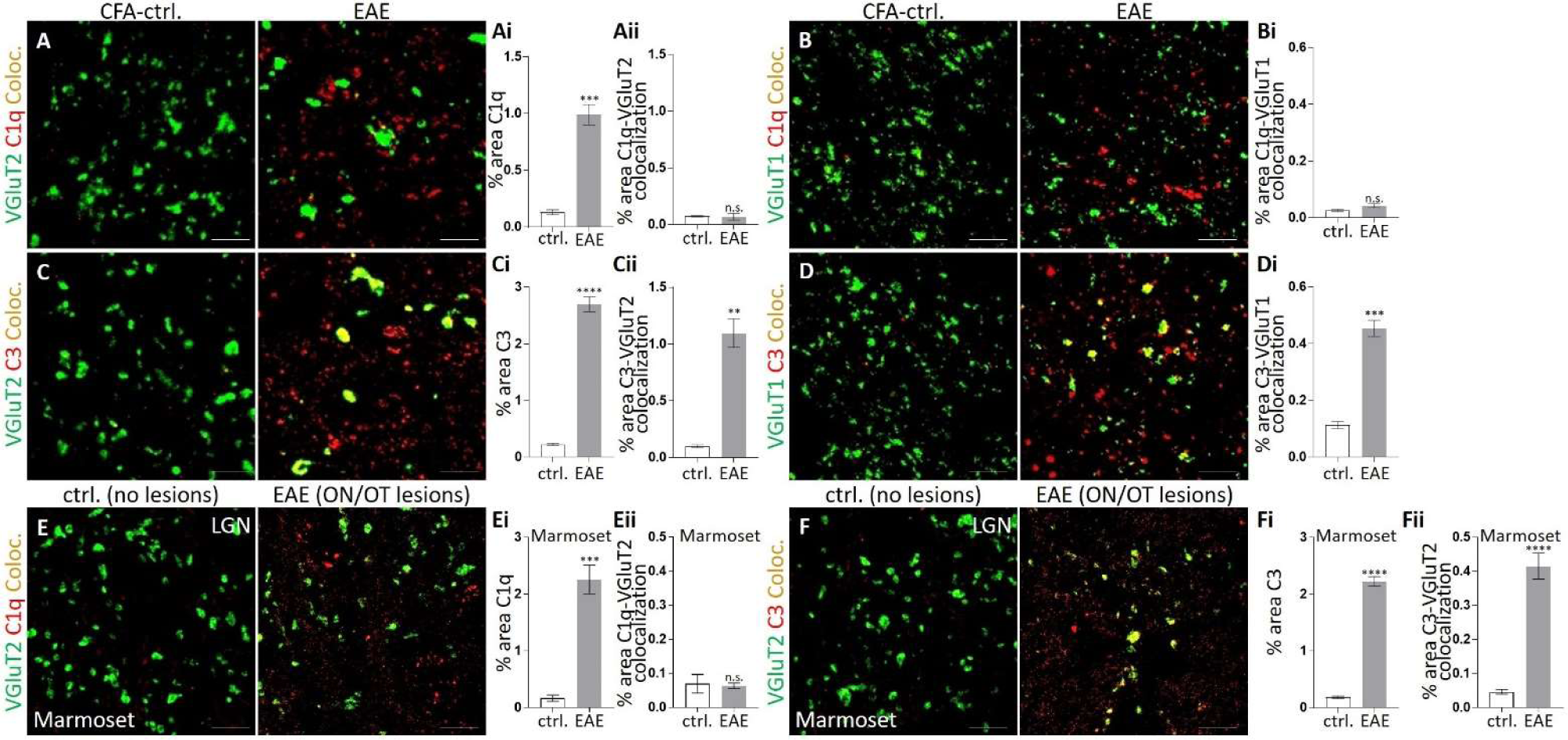
Complement component C3, but not C1q, localizes to synapses in mouse and marmoset EAE models. (**A,B**) Representative confocal images of control and EAE mouse LGN immunostained against presynaptic (**A**) VGluT2 (green) or (**B**) VGluT1 (green) and complement component C1q (red). While C1q is (**Ai**) significantly increased in the LGN in mouse EAE, (**Aii,Bi**) it does not localize to presynaptic compartments. (**C,D**) Representative confocal images from the LGN of control and EAE mouse LGN immunostained against presynaptic (**C**) VGluT2 (green) or (**D**) VGluT1 (green) and complement component C3 (red). Similar to C1q, C3 also (**Ci**) significantly increases in the LGN of EAE mice. Unlike C1q, C3 (**Cii,Di**) significantly colocalizes with presynaptic markers. (**A-D**) n=3. Scale bars, 5 μm. (**E,F**) Representative coronal sections of the LGN from non-EAE or EAE animals with no detectable lesions (control, no lesions) and marmosets that developed EAE with demyelinating lesions in the optic nerve and tract (EAE with ON/OT lesions) stained against presynaptic (**E,F**) VGluT2 (green) and (**E**) complement component C1q (red) or (**F**) complement factor C3 (red). Similar to mouse EAE, (**Ei**) C1q and (**Fi**) C3 increase in the LGN of marmosets that developed EAE with demyelinating lesions in the optic nerve and tract. Further quantification demonstrates significant colocalization of (**Fii**) C3, but not (**Eii**) C1q to VGluT2^+^ presynaptic compartments. (**E,F**) n=4. Scale bars, 5 μm. Data represent mean ± SEM, significant differences with **P < 0.01, ***P < 0.001, ****P < 0.0001, t-test. See also Fig. S6.

### AAV9-mediated expression of a C3 inhibitor at retinogeniculate synapses

To test whether synaptically localized, activated C3 was inducing microglia-mediated synapse engulfment and elimination, we took a gene therapy approach. We used adeno-associated virus 9 (AAV9) to overexpress the complement inhibitor Crry, which restricts C3 activation and inhibits C3-mediated opsonization, (Turnberg and Botto, 2003) in the retinogeniculate circuit. To gain further specificity, we fused Crry to a domain of complement receptor 2 (CR2), which is a membrane-associated receptor that binds activated C3. Following the sequence encoding the Crry-CR2-fusion protein was an autocleavage site and an EGFP sequence (Fig. 6A), which we used to identify transduced neurons and their projections. We termed this vector AAV-Crry. The expressed Crry-CR2 fusion protein is targeted by the CR2 domain to sites of activated C3 deposition where Crry then inhibits further C3 activation (Alawieh et al., 2015, Atkinson et al., 2005, Atkinson et al., 2006, Qiao et al., 2006, Alawieh and Tomlinson, 2016). For a control vector, we used an EGFP encoding construct (AAV-EGFP). AAV9 viral vectors were then generated and validated in cultured neuro-2a (N2a) cells. One week after viral transduction, western blot analysis revealed robust expression of Crry and EGFP in transduced N2a cells (Fig. 6B). Following initial validation, AAV-Crry or AAV-EGFPs were injected into the vitreous of both eyes of separate cohorts of four-week-old WT mice to transduce retinal ganglion cells (RGCs). Twenty-eight days following AAV delivery, EAE was induced in these mice (Fig. 6C). Mice were then analyzed at the onset of moderate clinical scores (average: AAV-Crry: 1.4±0.3; AAV-EGFP 1.5±0.3), which were typically observed around day 11 post-immunization (AAV-Crry: 11.2±0.6; AAV-EGFP: 11.4±0.4) (Fig. 7A). Using confocal imaging, we first determined the degree of EGFP colocalization with VGluT2^+^-retinogeniculate terminals in the LGN and found ∼45% of VGluT2^+^-retinogeniculate terminals in the LGN colocalized with EGFP after injection of both AAVs (Fig. 6D). We then immunostained for Crry protein and identified that Crry was highly enriched in presynaptic bouton structures in EGFP-labeled retinogeniculate arbors vs. other fine processes (i.e. axons) within the LGN of AAV-Crry injected mice (Fig. 6E). Previous work has demonstrated that these boutons represent VGluT2^+^-presynaptic terminals along retinogeniculate arbors (Hong et al., 2014), precisely where we originally identified enrichment of C3 (Fig. 5). These data demonstrate successful CR2-mediated Crry targeting to retinogeniculate synapses tagged by activated C3. In contrast, AAV-EGFP injected mice showed only diffuse Crry immunoreactivity with no concentration on or around retinogeniculate synapses in the LGN following EAE. Regardless of AAV treatment, mice showed comparable development of EAE clinical scores, infiltration of peripheral immune cells, and micro- and astrogliosis (Fig. 7A-D). These data suggest that retinogeniculate overexpression of Crry did not have global, systemic effects, which is important when considering the beneficial and detrimental effects of inflammation in the development of new therapeutic strategies.

**Figure 6:**
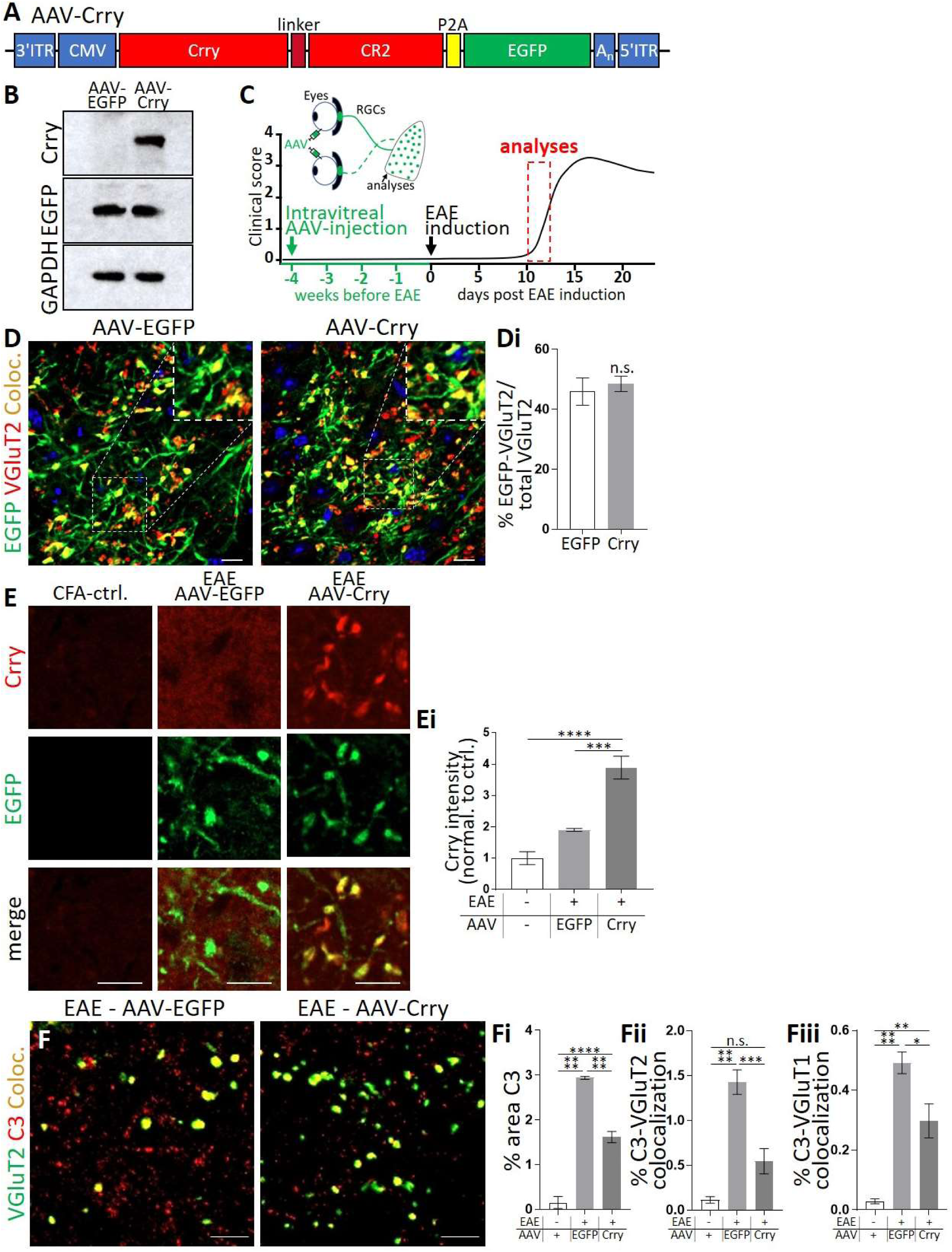
Creation and characterization of an AAV9 vector encoding the C3 inhibitor Crry. (**A**) Linear map of the AAV-Crry construct. ITR, inverted terminal repeat. CMV, human cytomegalovirus promotor. P2A, autocleavage site. A_n_, polyadenylation signal. (**B**) Representative immunoblots of Crry, EGFP, and GAPDH of homogenates from AAV-EGFP or AAV-Crry-EGFP (AAV-Crry) transduced neuro-2A cells reveals successful transduction of Crry in AAV-Crry treated cells compared to AAV-EGFP controls. (**C**) Timeline of *in vivo* AAV*-*rescue experiments. Mice received intravitreal AAV-injections to transduce retinal ganglion cells (RGCs) innervating the LGN 4 weeks prior to EAE induction. The LGN of mice was then analyzed at the onset of moderate clinical symptoms ∼11days post-immunization. (**D**) Representative confocal images of the LGN of AAV-EGFP or AAV-Crry-EGFP (AAV-Crry) injected mice stained against EGFP (green) and presynaptic VGluT2 (red). DAPI, nuclei counterstain (blue). Scale bars, 10 μm. Inset shows magnified details. (**Di**) Colocalization (Coloc., yellow) analysis reveals comparable localization of EGFP with VGluT2^+^-retinogeniculate terminals with both AAV-EGFP and AAV-Crry treated mice. (**E**) Representative confocal images of the LGN of control (injection of CFA without MOG_35-55_-peptide) and EAE-induced mice that received intravitreal injections with AAV-EGFPs or AAV-Crry. Immunostaining against Crry and EGFP reveals Crry is highly enriched at retinogeniculate presynaptic bouton structures labeled with EGFP following AAV-Crry-treatment. (**Ei**) Quantification of Crry levels (data normalized to CFA-controls) shows significantly increased Crry in the LGN in AAV-Crry-transduced EAE mice. (**F**) Representative confocal image of the LGN of AAV-EGFP (left) or AAV-Crry-treated (right) EAE mice immunostained against presynaptic VGluT2 (green) and complement factor C3 (red). (**Fi**) Overall C3 levels and (**Fii**) synapse-associated C3 are significantly reduced in the LGN of AAV-Crry-treated EAE mice. (**Fiii**) C3 is also reduced at VGluT1^+^-corticothalamic presynaptic inputs in AAV-Crry-treated EAE mice. (**E,F**) Scale bars, 5 μm. (**D-F**) n=5. Data represent mean ± SEM, significant differences with *P < 0.05, **P < 0.01, ***P < 0.001, ****P < 0.0001, (**C**) t-test, (**E,F**) ANOVA with Tukey’s *post hoc* test.

**Figure 7:**
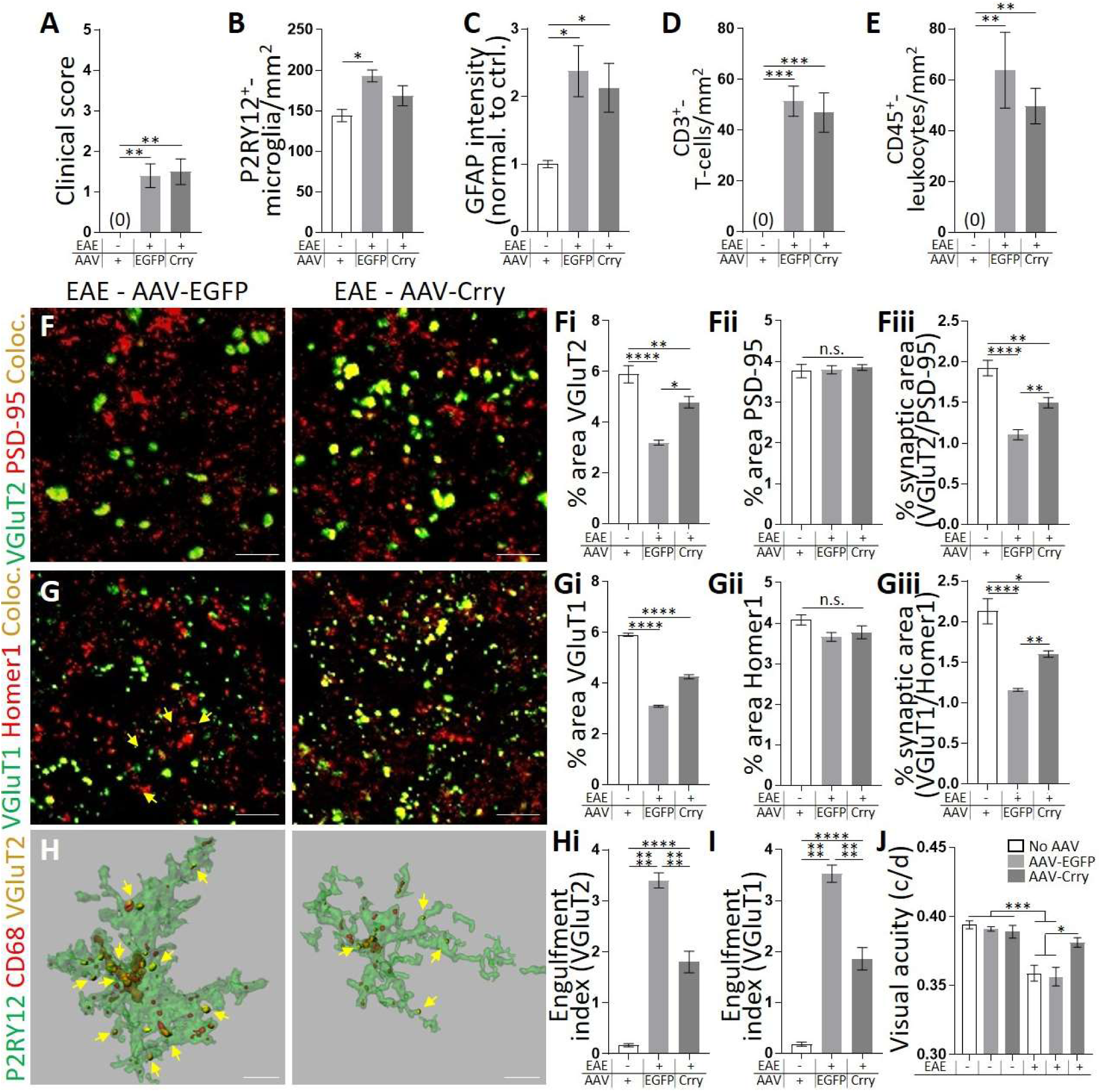
Overexpression of the C3 inhibitor Crry with AAV9 in the retinogeniculate circuit blocks microglial synaptic engulfment and synapse loss in the LGN and restores visual function. (**A**) Regardless of AAV treatment mice showed comparable development of clinical scores following EAE. (**B-E**) Within the LGN, we still detected (**B**) microgliosis as indicated by a reduction in P2RY12^+^-microglia, albeit no longer significantly different from controls. There was also still a significant increase in (**C**) GFAP^+^-astrocytes, (**D**) CD3^+^-T-cells and (**E**) CD45^+^-leukocytes following EAE in AAV-Crry treated mice. (**F,G**) Representative confocal images of the LGN of AAV-EGFP (left) and AAV-Crry treated (right) EAE mice immunostained against (**F**) presynaptic VGluT2 (green) and postsynaptic PSD-95 (red) or (**G**) presynaptic VGluT1 (green) and postsynaptic Homer1 (red). There is significant attenuation of (**Fi**) presynaptic VGluT2, (**Gi**) presynaptic VGluT1, and (**Fiii,Giii**) structural synapse (colocalized pre and postsynaptic markers, Coloc., yellow) loss in AAV-Crry treated EAE mice. Density of (**Fii,Gii**) postsynaptic markers is similar to controls in all conditions. (**H**) Representative 3D-surface rendering of a P2RY12^+^-microglia (green) and engulfed VGluT2^+^-retinogeniculate inputs (yellow) within CD68-labeled microglial lysosomes (red) in AAV-EGFP (left) and AAV-Crry treated (right) mice following EAE. (**H,I**) Engulfment of presynaptic inputs (arrows) in EAE is significantly attenuated with AAV-Crry treatment. (**J**) Visual acuity, measured as spatial frequency threshold (cycles/degree) in the optomotor test before the induction of EAE (three left columns) and at the onset of clinical symptoms (three right columns) of control and AAV-EGFP or AAV-Crry transduced mice. There is a significant decline in the visual acuity of non-transduced and AAV-EGFP transduced mice following EAE that is attenuated in AAV-Crry transduced mice. (**A,F-I**) n=5. (**B-D**) n=4. (**J**) n=5-8. Scale bars, (**F,G**) 5 μm, (**H**) 10 μm. Data represent mean ± SEM, significant differences with *P < 0.05, **P < 0.01, ***P < 0.001, ****P < 0.0001, ANOVA with Tukey’s *post hoc* test. See also Fig. S7.

### Crry-mediated inhibition of activated C3 at retinogeniculate synapses blocks microglial synapse engulfment, rescues synapse loss, and restores visual function

Following validation that Crry is successfully expressed using our AAV9 strategy and localizes to retinogeniculate presynaptic terminals, we asked if Crry overexpression reduced deposition of C3 following EAE. While we found modest reductions in C3 levels throughout the LGN in EAE AAV-Crry treated mice (Fig. 6F), there was a pronounced decrease in the colocalization of C3 with VGluT2^+^-retinogeniculate terminals in AAV-Crry treated EAE mice compared to AAV-EGFP controls. Interestingly, C3 colocalization with VGluT1^+^-corticothalamic inputs was also reduced, but this decrease was less pronounced compared to VGluT2^+^-terminals. Strikingly, the decrease in synaptic C3 deposition in AAV-Crry treated EAE mice was accompanied by a significant attenuation of VGluT2^+^- and VGluT1^+^-synapse loss following EAE (Fig. 7F,G). Consistent with microglia-mediated synapse loss via C3, there was also a significant reduction in VGluT2^+^- and VGluT1^+^-terminals engulfed within microglial lysosomes of AAV-Crry-treated mice compared to AAV-EGFP controls (Fig. 7H,I). To determine if these effects were specific to synapses within the LGN, we assessed engulfment and loss of presynaptic terminals in layer IV of the somatosensory cortex, a brain region without direct inputs from RGCs or the LGN. While microglial synaptic engulfment was detected and the density of presynaptic terminals was reduced following EAE, AAV-Crry did not protect against synapse engulfment or loss in this brain region (Fig. S7). This is consistent with the AAV-Crry injected mice also developing comparable EAE to AAV-EGFP controls. These data suggest that protective effects of AAV-mediated Crry expression are circuit-specific. This is particularly important to think about in the context inflammation, which can serve to resolve or exacerbate disease depending on the timing and context. For example, synapse loss may be beneficial in some cases to block excitotoxic damage at early stages of disease. Being able to protect specific synapses at specific times offers a more targeted strategy.

Finally, to ask if protection of synapse loss in the retinogeniculate circuit has consequences for visual function, we assessed visual acuity (Prusky et al., 2004) in mice before induction of EAE and at the onset of moderate clinical scores by optomotor testing ± AAV transduction (Fig. 7J). We first found that injection of the AAV intravitreally had no significant effect on visual acuity performance prior to EAE induction. We then observed a significant decline in visual acuity following EAE induction in all mice, except the AAV-Crry treated cohort. Strikingly, AAV-Crry treatment blocked loss of visual acuity, which was comparable to controls prior to EAE as well as previously published data assessing visual acuity in wild-type mice of a similar age (Schafer et al., 2016). These data demonstrate that AAV-Crry treatment can protect retinogeniculate circuit synapses and prevent functional visual impairment in demyelinating disease (Fig. 7J). We find C3 is enriched at synapses (Figs. 5; S6), there is no measurable neuronal degeneration within the retinogeniculate circuit, (Figs. 2; S4), and Crry is enriched in presynaptic compartments following AAV delivery (Fig. 6E). Therefore, Crry is likely mediating protection of synapses and visual function through local C3 inhibition at the synapse. Importantly, this is the first evidence in any context that inhibition of C3 at specific synapses may be possible using a gene therapy approach, which has significant therapeutic potential for many neurodegenerative diseases.

## Discussion

In postmortem human tissue, a preclinical nonhuman primate model of MS, and two murine models of demyelinating disease we have identified microglial engulfment of presynaptic inputs and synapse loss in a MS-relevant visual circuit, the retinogeniculate system. We further provide compelling evidence that this process can occur independent of demyelination, neuron degeneration, or peripheral immune infiltration. Instead, synapse engulfment and loss in the visual thalamus in all species is concomitant with local gliosis and occurs in the presence of synaptically enriched C3, but not synapse-associated C1q. Finally, overexpressing Crry, an inhibitor of C3, in the retinogeniculate circuit, results in a reduction in synaptic C3, inhibition of microglial synapse engulfment, attenuation of synapse loss specifically within the retinogeniculate system versus other brain regions, and protection of visual function. Together, these data support a model by which microglia mediate synapse loss through the alternative complement cascade in demyelinating disease, and this can occur independent of other myelin and neuronal pathology or peripheral immune cell infiltration. These data are the first to demonstrate that complement-mediated phagocytic signaling can modulate synaptic connectivity in demyelinating disease. We also provide a targeted gene therapy approach to inhibit activated C3 at synapse and protect structural and functional synapses with circuit-specificity – an approach that may be broadly applicable to many neurodegenerative diseases.

Our data in murine and nonhuman primate MS models suggest that microglia-mediated synapse engulfment and synapse loss occur prior to cell death and axonal degeneration. We provide further evidence in murine models that these events can also occur independent of demyelination and peripheral immune cell infiltration. This is in line with previous work suggesting that gray matter degeneration in MS can at least partially occur independent of demyelination (Mandolesi et al., 2015). Also, there is new evidence that, even with the new generation of FDA-approved MS therapies, there is ‘silent progression’ of the disease where a symptom-free patient at a 2 year follow-up still developed significant disability by the 10^th^ year (Cree et al., 2019). Local gliosis and microglia-mediated synapse elimination may be an underlying feature of this ‘silent progression’ and an early event in the progressive course of neurodegeneration. Consistent with this idea, prior work has implicated the thalamus as one of the earliest targets of degeneration in MS, and thalamic degeneration is an accurate predictor of subsequent MS-related disability (Eshaghi et al., 2018, Zivadinov et al., 2013). These data are also consistent with work showing synapse engulfment and early synapse loss in mouse models of other neurodegenerative diseases, where reactive microglia are also present (Hong et al., 2016, Paolicelli et al., 2017, Wishart et al., 2006, Yoshiyama et al., 2007). It is also important to consider the contribution of other immune and glial cells in this process. While our findings in the DTA mouse model suggest that complement-mediated microglial engulfment and synapse loss can occur in the absence of peripheral immune cell infiltration, there is still a possibility that these cells are involved in synapse loss in other contexts. For example, a recent study has shown that β-synuclein-specific T-cells, which are also present in human MS cases, can induce local gliosis and cortical gray matter degeneration in mice (Lodygin et al., 2019). In addition, other work has shown complement deposition is less in cortical regions compared to deep gray matter in MS (Brink et al., 2005). This raises the intriguing possibility that synapse loss may be regulated by T-cell-mediated mechanisms in cortical regions and complement-dependent innate immune mechanisms in deeper gray matter structures such as the thalamus.

It is also important to consider the role of astrocytes and potential astrocyte-microglia crosstalk. In development, similar to microglia, it has been shown that astrocytes can engulf synaptic material and regulate synaptic architecture (Chung et al., 2013). It has also been shown that reactive astrocytes become phagocytic and engulf increased amounts of synaptic material following stroke-induced brain ischemia and other disease-relevant conditions (Morizawa et al., 2017, Bellesi et al., 2017, Chung et al., 2016). In contrast, we show that synapse engulfment is mediated by microglia, but not astrocytes, at least in early stages of demyelinating disease. While this is consistent with reactive astrocytes showing decreased phagocytic capacity after LPS challenge (Liddelow et al., 2017), this does not rule out the possibility that local astrogliosis can influence microglia-mediated synapse elimination. Indeed, astrocytes have also been shown to promote microglial synaptic engulfment in the developing brain through IL-33 signaling (Vainchtein et al., 2018). Microglia-astrocyte crosstalk has also been explored in models of neurodegeneration where microglia have been shown to modulate astrocyte reactivity, transition to a neurotoxic A1 astrocyte phenotype, and astrocytic expression of C3 (Liddelow et al., 2017). These more reactive astrocytes could then, in turn, influence C3 levels at the synapses and/or the phagocytic or inflammatory state of microglia. The analysis of microglia-astrocyte crosstalk to disease progression and synapse loss will be important going forward.

Previous work in the mouse retinogeniculate system identified classical complement-dependent phagocytic signaling as a key mechanism regulating developmental engulfment and elimination of synapses by microglia (Schafer et al., 2012, Stevens et al., 2007, Bialas and Stevens, 2013). In these studies, C1q and C3 were localized to synapses. Microglia then engulf synapses, in part, through CR3, a receptor for C3. Similar microglia-mediated synapse elimination through the classical complement cascade has recently been identified in mouse models of Alzheimer’s disease, frontotemporal dementia and West Nile Virus infection (Hong et al., 2016, Vasek et al., 2016, Lui et al., 2016, Shi et al., 2017). Another recent study indicated that C1q and C3 are induced in MS and may colocalize with synaptic proteins in postmortem MS brains(Michailidou et al., 2015). However, it is unclear if this complement deposition is an independent pathological event or secondary to other brain pathology, and thus whether complement can induce synapse loss in demyelinating disease remained elusive. Our data indicate an increase in C1q, but no C1q deposition at synapses, in multiple MS models. Instead, we observe striking localization of C3 at synapses. These data suggest that synapse elimination can occur through the alternative, versus the C1q-dependent classical, complement cascade. These findings are in line with earlier work in EAE, suggesting that the alternative complement pathway predominates and that blocking this pathway globally prevents disease progression (Nataf et al., 2000). Similarly, clinical studies have shown increased plasma levels of C3 and elevated C3 in the CSF of MS patients, which correlate with disease severity and clinical disability (Ingram et al., 2012, Ingram et al., 2014, Tatomir et al., 2017). Interestingly, patients with progressive MS, particularly those with primary progressive MS, showed the highest concentrations of C3 in the CSF and plasma (Ingram et al., 2012, Aeinehband et al., 2015).

Despite compelling evidence for the alternative cascade in MS, it is still important to consider the contribution of C1q. For example, our data demonstrate elevated levels of C1q, and it has been shown recently that microglial-derived C1q can increase astrocyte reactivity and reactive astrocytes production of C3 (Liddelow et al., 2017). This astrocyte-derived C3 could then localize to synapses. It is unknown what cell types are producing C3 and C1q in our demyelinating models, which will be an important future direction. Related, it is unknown what cognate receptor the microglia are using to phagocytose C3-opsonized synapses in our models. Previous work identified an ∼50% block of microglial synaptic engulfment in mice deficient in complement receptor 3 (CR3), during developmental synaptic pruning and in a mouse model of Alzheimer’s disease (Hong et al., 2016, Schafer et al., 2012). Another study showed that West Nile Virus-induced synapse loss and cognitive impairment involves complement receptor 3a (C3aR), but not CR3 (Vasek et al., 2016). However, assessing synapse loss CR3 or C3aR-deficient mice is challenging, as both mouse lines show attenuated EAE due to a dampened peripheral inflammatory response (Boos et al., 2004, Bullard et al., 2005). Therefore, the identification of microglial receptors involved in synapse engulfment in demyelinating disease requires new genetic tool development to more specifically manipulate these receptors in microglia versus other myeloid lineage cells. Elucidating the microglial receptor could also be an interesting direction in light of recent work showing that fibrinogen leakage from the vasculature in the mouse EAE model of MS can induce clustering of microglia around vascular networks and axon loss via CR3 (i.e. Cd11b) (Davalos et al., 2012). A follow-up study in an Alzheimer’s model further demonstrates that this fibrinogen-CR3 interaction could also modulate dendritic spine loss through modulation of microglial release of reactive oxygen species (Merlini et al., 2019). Exploring a potential link between blood brain barrier disruption, fibrinogen leakage, and microglial receptors required for synapse elimination in MS could be another interesting future direction.

Our data demonstrate that synapse loss can occur independent of other MS-relevant pathology and that C3 is highly enriched at synapses vs. other neuronal compartments at early stages of EAE. Further, using an AAV-mediated *in vivo* gene delivery approach, we show that when the C3 inhibitor Crry is fused to a domain of CR2, the receptor for activated C3, Crry is highly enriched in presynaptic boutons. Most importantly, C3 inhibition with Crry can protect synapses and visual function. Taken together, we provide an exciting new therapeutic strategy to prevent synapse loss and visual impairment within specific CNS circuits in neurodegenerative disease. Supporting this strategy as a viable therapeutic option, AAV-mediated delivery of Crry has been utilized to decrease C3 activation in models of macular degeneration and ischemic brain injury (Atkinson et al., 2005, Huang et al., 2008, Marshall et al., 2014, Alawieh et al., 2015, Atkinson et al., 2006). Another study relevant to MS showed that systemic, peripheral administration of a Crry-CR2 encoding AAV attenuated EAE (Hu et al., 2012), which was attributed to the targeting of peripheral immune cells and restricting epitope spreading. This is in contrast to our study, which is focused on the neurodegenerative aspects of MS in a specific, deep gray matter brain circuit. During neurodegeneration, inflammation can be beneficial as well as detrimental depending on the stage of disease and CNS region affected. For example, synapse loss may be a compensatory mechanism in some disease contexts to block excitotoxicity. However, these mechanisms of immune-mediated synapse loss may become dysregulated over the course of disease leading to aberrant elimination of other ‘innocent bystander’ synapses. Also, synapse loss within some circuits or diseases may occur independent of complement (Brink et al., 2005, Di Liberto et al., 2018, Lobsiger et al., 2013). Thus, this targeted, circuit-specific therapy could offer significant therapeutic potential to target specific circuits and not others during neurodegeneration. Further supporting our strategy as a viable therapeutic option in disorders involving retinal cells and associated brain circuits, the FDA recently approved an AAV based gene therapy approach to treat rare forms of inherited vison loss via subretinal AAV-injections (Russell et al., 2017). Other neurodegenerative diseases or synapse loss in other CNS regions in MS could also benefit from such a strategy, but this would likely require systemic delivery and/or cell-type specific promotors to target specific CNS neuron populations. Together, our data support AAV delivery of C3 inhibitors fused to the activated C3 receptor CR2 as a viable therapeutic approach to block synapse loss at specific neural circuits in neurodegeneration.

In summary, we provide compelling evidence that local reactive microglia engulf and eliminate synapses in a manner dependent on synaptically localized complement factor C3. This includes data from human MS tissue that shows significant synapse loss and microglial engulfment of synapses. We further provide evidence from multiple complementary MS models that synapse loss can occur independent of demyelination, neuron degeneration, and peripheral immune cell infiltration. In contrast, synapse loss can be a separable degenerative process that is positively correlated with local gliosis. With our novel AAV approach, we provide the first evidence that targeted inhibition of C3 at synapses is sufficient to block synapse engulfment and elimination by microglia and can protect from visual function decline. With the growing number of neurological diseases that appear to involve complement and microglia in synapse pathology, these data offer a viable therapeutic target to protect synapse integrity and function in disease.

## Acknowledgments

We thank Dr. Christopher C. Hemond (UMMS) for critical reading of the manuscript. We thank Dr. Paola Perrat (UMMS) for advice on the design and production of plasmids, Shannon Becker (UMMS) and Anoushka Lotun (UMMS) for assistance with tissue preparation, and Dr. Claudio Punzo and Georgia Gunner for advice and training on optomotor testing. We further thank Prof. Dr. Matthew Rasband (Baylor College of Medicine) for sharing CASPR and βIV-spectrin specific antibodies and Prof. Oleg Butkovsky for providing P2RY12 specific antibodies. This work was supported by DFG (Deutsche Forschungsgemeinschaft) grant WE 6170/1-1 (SW), NIMH – R00MH102351 (DPS), NIMH – RO1MH113743 (DPS), NIMH – R21MH115353 (DPS), Charles H. Hood Foundation (DPS), Brain & Behavior Research Foundation (DPS), Worcester Foundation (DPS), the Intramural Research Program of NINDS (DSR), and the Dr. Miriam and Sheldon G. Adelson Medical Research Foundation (DPS, DSR, BP).

## Author Contributions

S.W. and D.P.S. designed the study and wrote the manuscript. S.W. performed and analyzed the experiments. J.J. assisted with mouse maintenance and conduction and analysis of experiments. S.J.C. and C.M.W. provided first murine EAE tissue and assisted in establishing the EAE mouse model. G.G. assisted with viral vector production. B.P. and R.B.K. maintained DTA mice and collected tissue for analyses. D.S.R. supervised collection and processing of human brain tissue by S.K.H. and treatment and collection of marmoset tissue by N.J.L. All authors revised the manuscript for intellectually important content.

## Declaration of Interests

The authors declare no conflict of interest.

## STAR Methods

### Animals

Wildtype C57Bl/6J mice (stock #000664) were obtained from Jackson Laboratories (Bar Harbor, ME). *Plp1-CreER*^*T*^;*ROSA26-EGFP-DTA* (DTA) and *ROSA26-EGFP-DTA* (DTA-ctrl.) mice were generated as previously described (Traka et al., 2010, Traka et al., 2016). Littermate controls were used for all mouse experiments. 12 common marmosets (*Callithrix jacchus*), eight females and four males, were selected from the NINDS colony. All animal experiments were performed in accordance with Animal Care and Use Committees (IACUC) and under NIH guidelines for proper animal welfare.

### Experimental autoimmune encephalomyelitis induction in mice

As described previously (Crocker et al., 2006), experimental autoimmune encephalomyelitis (EAE) was induced in 8-week-old male mice, by subcutaneous (s.c.) administration of 200 μg of MOG_35-55_ peptide (AnaSpec Inc., AS-60130-10, Fremont, CA, USA) emulsified in complete Freund’s Adjuvant (CFA, Sigma, F5881, Saint Louise, MO, USA) containing 0.2 mg of *Mycobacterium tuberculosis* H37Ra (Difco Laboratories, #231141 Detroit, MI, USA) into the flanks of both hind-limbs. Control animals received s.c. injections lacking MOG_35-55_ peptide. At the time of immunization and 48h later, mice further received intraperitoneal (i.p.) injections with 500 ng of pertussis toxin (List Biologicals, #180, Deisenhofen, Germany). Weights and clinical scores were recorded daily (score 0.5: distal tail limpness; score 1: complete limp tail; score 1.5: limp tail and hindlimb weakness; score 2: mild hindlimb paresis; score 2.5: unilateral hindlimb paralysis; score 3: bilateral hindlimb paralysis, score 4: moribund). For the EAE experiments presented in this study, mice displaying moderate clinical scores were analyzed at the onset of EAE which was typically observed between day 10-12 post immunization. None of the CFA-treated control mice developed clinical symptoms.

### EAE induction in marmosets

EAE was induced in marmosets by injecting human white matter homogenate emulsified in complete Freund’s adjuvant (Difco Laboratories, BD, Franklin Lakes, NJ, USA) (Absinta et al., 2016, Lee et al., 2018). For EAE induction, intradermal injections were divided over four areas around the inguinal and axillary lymph nodes. To conserve animals, marmoset tissue used in this study was drawn from prior studies, some of which were designed to examine the effects of intranasal inoculation of human herpesvirus (HHV)-6 on EAE development. Therefore, a subset of marmosets (EAE and non-EAE) used in our study also received either HHV6 serotype A derived from SUP-T1 cells, HHV6 serotype B derived from HSB-2 cells, or were inoculated with uninfected control SUP-T1 cells. Importantly, although the onset of EAE was accelerated with HHV6 treatment, no pathological changes in or outside of lesions other than accumulation of viral material were observed in the HHV6 studies (Leibovitch et al., 2018). Marmoset demyelinating lesions were monitored in vivo and ex vivo (i.e. postmortem dissected brain) by MRI. The animals were subsequently divided into 2 groups. The experimental group included EAE marmosets with clear demyelinating lesions in the optic nerve and tract. The control group were either EAE or non-EAE marmosets with no evidence of demyelinating lesions in the optic nerve or tract.

### Tamoxifen injections of DTA mice

Male *Plp1-CreER*^*T*^;*ROSA26-EGFP-DTA* mice 5-7-week-old were injected intraperitoneally with 0.8 mg of 4-hydroxytamoxifen (Sigma) per day for four consecutive days, as previously described, to induce demyelination driven by genetic ablation of mature oligodendrocytes (Traka et al., 2010, Traka et al., 2016). Tamoxifen-injected ROSA26-EGFP-DTA littermates were used as control mice.

### Immunostaining of mouse tissue

At the indicated time points, mice were deeply anesthetized and transcardially perfused with 0.1M phosphate buffer (PB) followed by 4% paraformaldehyde (PFA)/0.1M PB. Brains, retinas, and optic nerves were dissected, and brains and retinas were post-fixed at 4°C in PFA for 4h and 30 min, respectively. Brains and optic nerves were then equilibrated in 30% sucrose/0.1M PB, and then embedded in a 1:1 mixture of 30% sucrose/0.1M PB and O.C.T. compound (ThermoFisher Scientific Waltham, MA, USA). Tissue was cryo-sectioned into 10 μm coronal brain sections and longitudinal optic nerve sections. Sections and whole retinas were blocked and permeabilized at room temperature for 1h in 10% normal goat serum/0.1M PB containing 0.3% Triton-X 100 (all Sigma) followed by overnight incubation with primary antibodies at room temperature. The following primary monoclonal (mAb) and polyclonal (pAb) antibodies have been used: mouse mAb α-ALDH1L1 (clone N103/39, Millipore, MABN495, 1:1000), mouse mAb α-APP (clone 22C11, Millipore, MAB348, 1:200), rabbit mAb α-C1q (clone 4.8, Abcam, ab182451, 1:100), rat mAb α-C3 (clone 11H-9, Abcam, ab11862, 1:500), rabbit pAb α-CASPR (provided by Matthew N. Rasband, 1:100), rabbit mAb α-CD3 (clone SP7, Abcam, ab16669,1:100), rat mAb α-CD45 (clone IBL-3/16, BioRad, MCA1388, 1:100), rat mAb α-CD68 (clone FA-11, AbD Serotec, MCA1957, 1:1000), rabbit pAb α-cleaved caspase-3 (Cell Signaling Technologies, #9661, 1:200), mouse mAb α-Crry (clone TLD-1C11, Santa Cruz, sc-53530, 1:100), rabbit pAb α-Factor H (Abcam, ab170036, 1:100), mouse mAb α-GFAP (clone clone G-A-5, Sigma, G3893, 1:500), chicken pAb α-EGFP (Abcam, ab13970, 1:500),rabbit pAb α-Homer1 (Synaptic Systems, #160003, 1:1000), rabbit pAb α-Iba1 (Wako Chemicals, #019-19741, 1:500), rat mAb α-LAMP2 (clone GL2A7, Abcam, ab13524, 1:200), mouse mAb α-MAG (clone 513, Millipore, MAB1567, 1:100), rat mAb α-MBP (clone 12, Millipore, MAB386, 1:500), mouse mAb α-MOG (clone 8-18C5, Millipore MAB5680, 1:200), rabbit pAb α-Neurofilament 200 (Sigma, N4142, 1:1000), chicken pAb α-NeuN (Millipore, ABN91, 1:1000), rat mAb α-P2RY12 (clone S16007D, BioLegend, 848002, 1:100), mouse mAb α-PSD-95 (clone 6G6-1C9, Millipore, MAB1596, 1:100), rabbit pAb α-βIV-spectrin provided by Matthew N. Rasband, 1:100), guinea pig pAb α-VGluT1 (Millipore, ab5905, 1:2000), and guinea pig pAb α-VGluT2 (Millipore, ab2251, 1:2000). The following day, sections were incubated with appropriate Alexa-fluorophore-conjugated secondary antibodies (ThermoFisher Scientific) and mounted with vectashield containing DAPI (Vector laboratories, Burlingame, CA, USA).

### Collection and immunostaining of human and marmoset tissue

Human brain tissue was dissected and formalin-fixed at autopsy after obtaining informed consent for collection. Tissue collection followed protocols approved by NIH Institutional Review Board. Marmoset brains were collected within 1h of death and fixed in 4% paraformaldehyde. Once individual slabs containing the lateral geniculate nucleus (LGN) of human and marmoset brains were identified, tissue was embedded in paraffin and sliced into 10 μm thick slides. For immunohistological stains on marmoset and human tissue, slices were deparaffinized and rehydrated, and antigen-binding sites were retrieved by heating for 1h in 10 mM citrate buffer, pH 6.0, with 0.05% Tween-20 in a steam oven before staining. All subsequent steps followed the above described protocol for staining of mouse tissue. If goat-derived primary antibodies were used, blocking was performed with 10% donkey serum (Sigma). Different from staining of mouse tissue, rabbit pAb α-C1q (Novusbio, NBP1-87492, 1:200) and mouse mAb α-C3d (clone 7C10, Abcam, ab17453, 1:50) were used to detect complement components, chicken pAb α-MAP2 (EnCor Biotechnology, CPCA-MAP2, 1:1000) was used to stain neurons, and rabbit pAb α-VGluT1 (Millipore, abN1627) was used to label presynaptic corticothalamic inputs and in addition to the antibodies used on mouse tissue goat pAb α-Iba1 (Abcam, ab5076, 1:200) was used to visualize microglia on human and marmoset tissue.

### Engulfment analysis

Engulfment analysis was performed according to previously described protocols (Schafer et al., 2012, Schafer et al., 2014). Briefly, two sections from each sample containing the LGN in marmoset and human and the dorsal LGN in mouse were immunostained and imaged on a Zeiss Observer Spinning Disk Confocal microscope equipped with diode lasers (405 nm, 488 nm, 594 nm, 647 nm) and Zen Blue acquisition software (Zeiss; Oberkochen, Germany). For each hemisphere, 2-3 randomly chosen 63x fields of view within the dorsal lateral geniculate nucleus were acquired with 50-70 z-stack steps at 0.22 μm optimal spacing using identical settings. Images were then processed in ImageJ (NIH) and individual images of 15-20 single cells per animal were processed in Imaris (Bitplane; Zurich, Switzerland) as previously described (Schafer et al., 2014, Schafer et al., 2012). Engulfment analysis was restricted to synaptic material within CD68-positive microglial lysosomes or LAMP2-positive astrocytic lysosomes. Unbiased quantification of all images was performed blind to genotype or treatment of animals.

### Synapse and complement density analyses

For assessing the density of presynaptic inputs (VGluT1, VGluT2) and postsynaptic compartments (Homer1, PSD-95), and for determining the deposition of complement components (C1q, C3), two stained sections from each sample containing the LGN (human and marmoset) or dorsal LGN (mice) were imaged on a Zeiss LSM700 laser scanning confocal microscope equipped with 405 nm, 488 nm, 555 nm, and 639 nm lasers and Zen black acquisition software (Zeiss; Oberkochen, Germany). For each hemisphere, 2-3 randomly chosen 63x fields of view within the LGN were acquired with three z-stack steps at 0.68 μm spacing. Identical settings were used to acquire images from all samples within one experiment, and data analyses were performed using ImageJ (NIH, version 1.52k) as described previously with minor modifications (Schafer et al., 2012, Hong et al., 2016). First, to determine a consistent threshold range, sample images for each genotype and condition were subjected to background subtraction and then manual thresholding blinded to condition and genotype for each channel within one experiment was performed (IsoData segmentation method, 85-255). Then, each channel from single z-planes of the z-stacks (3 z-planes per animal) were subjected to the same background subtraction and thresholding, which was kept consistent for a given experiment. Using the analyze particles function, the total area of presynaptic inputs, postsynaptic compartments and complement component deposition was measured from the thresholded images. To quantify the total area of structural synapses, the image calculator tool was first used to visualize co-localized pre- and postsynaptic puncta from the previously thresholded images. Then, the analyze particles function was used to calculate the total area of co-localized signals. Colocalization of complement components (C1q, C3) or Crry-EGFP with presynaptic puncta was performed similarly. Data from single planes was averaged for each z-stack of each field of view, and the mean of all fields of view from one animal was determined to calculate densities.

### Density analysis of myelin, nodes, paranodes and axons

Similar to the quantification of synapses, densities of myelin (MOG, MAG, MBP), axons (neurofilament, MAP2), as well as nodes of Ranvier (βIV-spectrin) and paranodal junctions (CASPR) were determined. In brief, 2-3 randomly chosen 20x fields of view within the region of interest (ROI; LGN, optic nerve, optic tract) of 2-3 slides per animal were acquired with three z-stack steps at 0.44 μm spacing from each animal on a Zeiss Observer Spinning Disk Confocal microscope. Identical settings were used for the acquisition of all images from one experiment, and areas outlining the desired ROI (LGN, retina, optic nerve) were selected and quantified blind to genotype or condition with ImageJ as described above for the synapse analyses.

### Cell density quantification

For determining the density of microglia (P2RY12), neurons (NeuN, MAP2), infiltrating peripheral immune cells (CD3, CD45), apoptotic cells (cleaved caspase 3) and degenerating axons (APP), single plane 10x and 20x fluorescence images were collected from both hemispheres of at least two slices from each animal with a Zeiss Observer microscope (Zeiss; Oberkochen, Germany). Identical settings were used for the acquisition of all images from one experiment, and areas outlining the desired ROIs (LGN, retina, optic nerve) were manually counted blind to treatment or genotype using Zen Blue software (Zeiss). The counts were normalized to the total area of the selected ROIs to calculate cell densities.

### Fluorescence intensity analysis

For determination of GFAP, P2RY12, and Crry fluorescence intensities in the LGN, 10x and 20x single plane epifluorescence images were collected from both hemispheres of at least two slices from each animal with a Zeiss Observer microscope equipped with Zen Blue software (Zeiss; Oberkochen, Germany). Identical settings were used to acquire images for all samples within one experiment and data analyses were performed using ImageJ (NIH, version 1.52k) and blinded to genotype or conditions of samples. Before quantification, thresholds of pixel intensity were set to the full range of 16-bit images to ensure a consistent pixel range across all images and background was subtracted from all images. To sample fluorescence intensity, a ROI covering the LGN was manually selected for each image and the raw integrated density of pixels within each ROI was measured. Average intensity over all ROIs was quantified for each animal and then normalized to the values of control treated animals.

### Generation of adeno-associated viral vector expressing Crry

The pAAV-CB6-PI plasmid (4045 bp) (Gene Therapy Center, UMass Medical School, Worcester, MA, USA) was used as vector backbone in this study. Sequences encoding the complement inhibitor Crry fused to a CR2 sequence and followed by an autocleavage side and the sequence for EGFP were synthesized as two separate gBlock gene fragments (IDT, Skokie, IL). The first gBlock (1382 bp) encoded the sequence for the mature murine Crry protein (residues 1-319, GenBank accession number NM013499) flanked by overhangs (∼25bp) with the backbone at the 5’-terminus and the second gBlock at the 3’-terminus (covering parts of the (G_4_S)_2_ linker sequence). The second gBlock (1645 bp) encoded the sequence for a (G_4_S)_2_ linker directly fused to a sequence encoding 4 N-terminal short consensus repeats of a CR2-sequence (residues 1-257 of mature protein, GenBank accession number M35684), followed by the sequence of a porcine teschovirus-1 2A (P2A) autocleavage site and the sequence for EGFP (New England Biolabs, NEBuilder Assembly Tool 2.0.8). At the 3’-terminus, the second gBlock was flanked by an overhang with the backbone and at the 5’-terminus with an overhang (both ∼25bp) identical to terminal parts of the Crry sequence of the 3’-end of the first gBlock. Both gBlocks were PCR-amplified, purified and cleaned before sequences were confirmed by Sanger sequencing. To assemble gBlocks with the backbone, the one-step isothermal NEBuilder HiFi DNA assembly cloning method has been utilized according to the manufacture’s recommendations (New England Biolabs, Ipswich, MA, #E2621L). 4 μl of the assembly reaction were used for transformation of Stbl3 chemically competent *E. coli* (Thermo Fisher Scientific, # C737303) and bacterial plasmid DNA extraction was performed using QIAfilter Plasmid Mega Kit (Qiagen, #12281) according to the manufacturer’s recommendations. Control constructs encoding the sequence for EGFP but, lacking the sequences for Crry fused to CR2, were generated by restriction enzyme digest (NheI/StyI) and religation of final construct. Restriction analysis of the constructs was performed with the following enzymes or combination of enzymes: SmoI, HincIII, XhoI/XbaI, BamHI, SalI/XhoI, XhoI, BamsHIEcoRV and HindIII (all from New England Biolabs). The final constructs were further verified by sequencing analysis and subsequently packaged as AAV9 vectors by the UMass Viral Vector Core of the Gene Therapy Center at the University of Massachusetts Medical School.

### Intravitreal injection of AAVs

Four-week-old mice were deeply anesthetized with isoflurane, the surrounding tissue around the eye was swabbed with 5% povidone-iodine, and an incision was made into the sclera (app. 2 mm posterior of the superior limbus) using a sterile, sharp 30G needle. After careful removal of the 30G needle, a 33G blunt needle (Hamilton, 1.5”/PT3, #7803-05) attached to a 10 μl gastight syringe (Hamilton, #1701) was carefully inserted into the same incision and viral suspension was injected into the vitreous without damaging the lens or retina. Viral suspension (3 μl) was injected at a titer of 2.5 ×10^13^ genome copies (GC)/ml. After injection, the needle was kept inside the vitreous for ∼1 min. Mice received injections into both eyes. Finally, ophthalmic antibiotic ointment and local anesthetic were applied. Mice were placed in their home cage on a 37°C heating pad and monitored until they recovered from anesthesia. Twenty-eight days after AAV-injection, EAE was induced in the mice as described above. Only mice that showed substantial colocalization (<35%) of EGFP with VGluT2^+^-retinogeniculate terminals in the LGN were included in the analyses.

### Culture and treatment of neuro-2a cells

Murine neuro-2a cells (N2a) were purchased from ATCC (#CCL-131) and cultured in DMEM (Thermo Fisher) supplemented with 10% fetal bovine serum (HyClone, USA) and 1% penicillin-streptomycin at 37°C in a humidified incubator. At ∼80% confluency, cells were passaged. For AAV-mediated transduction, N2a cells were seeded and, after full attachment to the plate, treated with 25 μM all trans-retinoic acid (RA, Sigma, Cat. R2625) to induce neurite outgrowth. The next day, cells were inoculated with either AAV-Crry or AAV-EGFP with a MOI of 10^5^ GC/cell. AAVs were removed 48h after infection and cells were harvested or fixed for analysis another five days later.

### Western blot

Samples were lysed in T-PER (Thermo Scientific), boiled at 95°C for 10 min, separated on 4-20% Mini-PROTEAN TGX Gels under reducing conditions, and transferred to polyvinylidene diflouride membrane. After blocking with 5% (w/v) non-fat milk/PBS (all from BioRad), membranes were incubated with the following antibodies: mouse mAb α-Crry (clone TLD-1C11, Santa Cruz, sc-53530, 1:100), rabbit pAb α-EGFP (Millipore, ab3080P, 1:1000), goat pAb α-GAPDH (Abcam, ab9483, 1:1000), and respective peroxidase-conjugated secondary antibodies (BioRad). Signals were visualized by enhanced chemiluminescence. Prior to the detection of further antigens on the same membrane, antibodies were washed off with Restore Plus Western Blot Stripping Buffer (Thermo Scientific).

### Optomotor testing

Assessment of visual acuity was performed before the induction of EAE and at the onset of clinical symptoms in awake, freely moving mice. Individual mice were placed on a round, elevated platform in the center of a soundproof chamber that was surrounded by four computer screens on which visual stimuli were projected. Mice were exposed to a virtual cylinder consisting of rotating sine wave gratings (12 deg/sec) of various spatial frequencies (cycles/degree). Mice reflexively track these gratings by stereotypic head movements as long as the grating is visible (Prusky et al., 2004). To determine grating acuity, first a homogeneous gray stimulus was projected on the cylinder, which was followed by a low-spatial-frequency (0.05 cyc/deg) sine wave grating of the same mean luminance that was randomly rotating in one horizontal direction. A video camera positioned in the lid of the chamber directly above the animal was used by a trained observer to record the mice and assess smooth, reflexive head movements in response to the rotating gratings. To determine the highest spatial frequency perceptible for each individual mouse, the spatial frequency of the grating was systematically increased until the animal no longer responded to the projected grating. Visual acuity was assessed blind to treatment of animals.

### Statistical analysis

Results are expressed as means ± standard error (SEM) from at least three independent replicates for each experimental group. Two-tailed Student’s t-test to compare two groups or one-way analysis of variance (ANOVA) followed by Tukey’s multiple comparison *post hoc* test to compare multiple groups was performed using Prism 7 software (Graph-Pad, La Jolla, CA, USA). Values of *P□<□0.05, **P□<□0.01, ***P□<□0.001, ****P□< □0.0001 were considered statistically significant.

