## Supplement Figures and Tables for "Targeted complement inhibition at synapses prevents microglial synaptic engulfment and synapse loss in demyelinating disease"

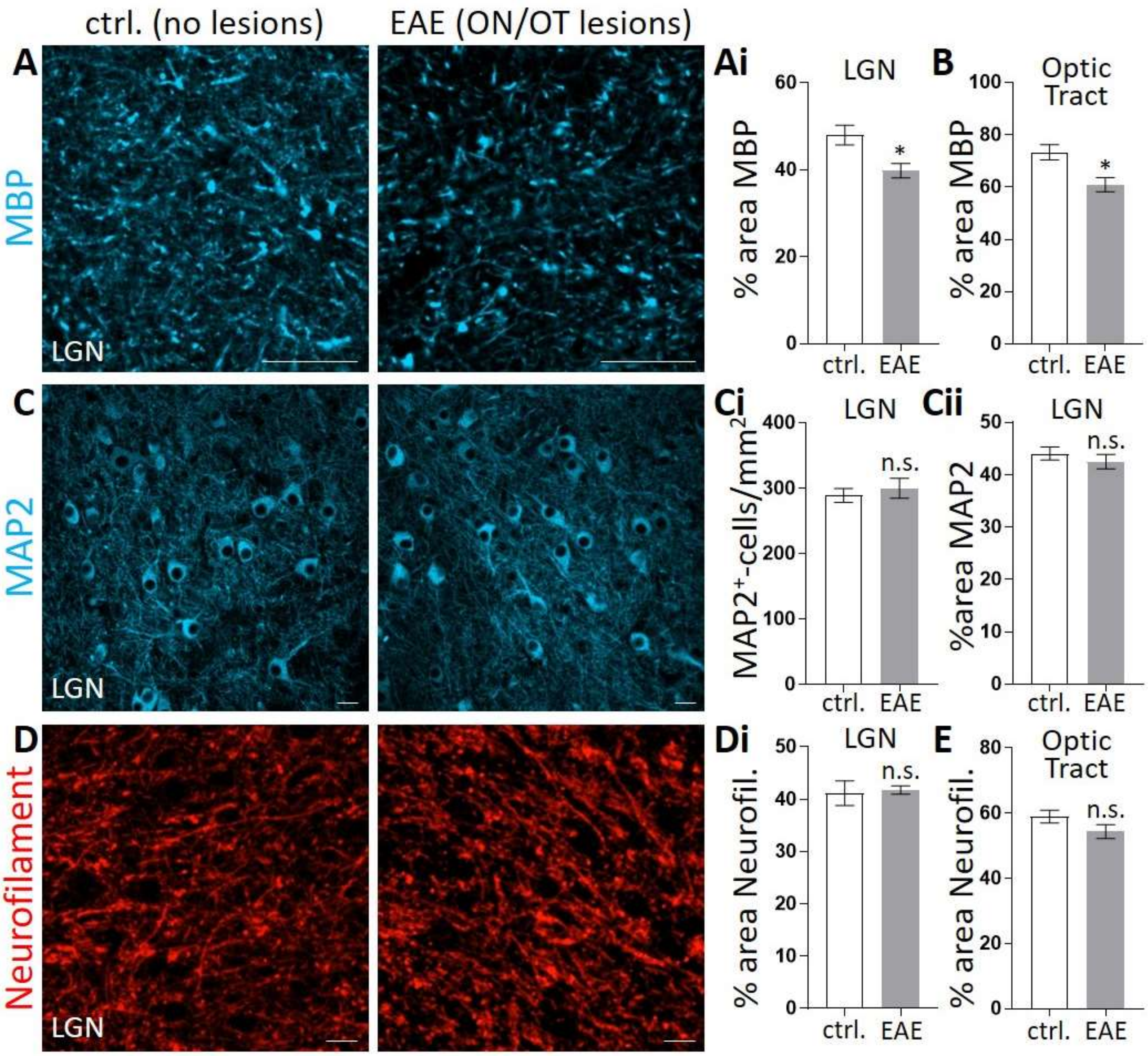


**Figure S1: Demyelination, but no significant cell death or axon degeneration, is detected in the LGN of the marmoset EAE model.** (**A**) Representative images of adjacent sections from (**A-E**) the same LGN non-EAE or EAE animals with no detectable lesions (control, no lesions) and marmosets that developed EAE with demyelinating lesions in the optic nerve and tract (EAE with ON/OT lesions) shown in Fig. 1 immunostained for myelin basic protein (MBP). (**Ai,B**) Quantification reveals moderate demyelination in the (**Ai**) LGN and (**B**) optic tract innervating the LGN of EAE-induced marmosets that developed ON/OT lesions. (**C-E**) Immunostaining against (**C**) MAP2 and (**D**) neurofilament (Neurofil.) in the LGN. There is no significant change in the overall (**Ci**) number or (**Cii**) density of $MAP2^{+}$-neurons in the LGN of EAE marmoset with ON/OT lesions compared to controls. There is also no significant decrease in the area of $neurofilament^{+}$-axons in the (**Di**) LGN and (**E**) optic tract of marmosets that developed EAE with ON/OT lesions. (**A-E**) n=6 marmosets. Scale bars, 20 μm. Data represent mean ± SEM, significant differences with *$P < 0.05$, t-test.

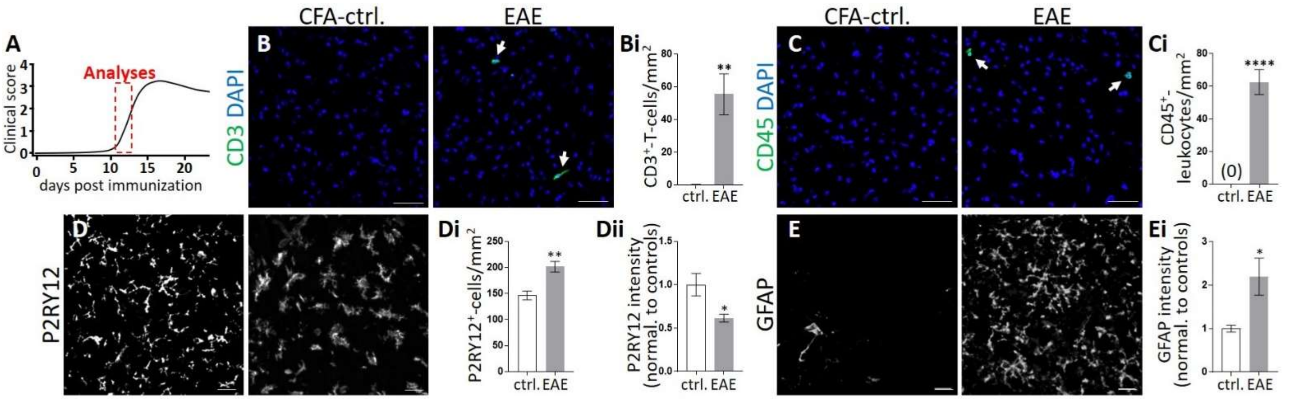


**Figure S2: Peripheral immune cell infiltration, microgliosis and astrogliosis in the LGN at early stages of EAE in the mouse.** (**A**) Schematic of the course of clinical symptoms in EAE of C57Bl6/J WT mice. Mice were analyzed at the onset of moderate clinical symptoms, typically observed between day 10-12 post-immunization. (**B-E**) Note, all immunostaining was performed in adjacent sections from the same LGN tissue analyzed in Figs. 2-3. (**B,C**) Immunostaining against (**B**) CD3 (green) and (**C**) CD45 (green) to assess infiltration of peripheral immune cells (arrows) in LGN from EAE mice 10-12 days post-immunization. DAPI was used to label nuclei (blue). There is a significant increase in (**Bi**) $CD3^+$-T-cells and (**Ci**) $CD45^+$-leukocytes in the LGN from EAE mice compared to CFA-treated controls. (**D**) Coronal sections of the LGN stained against P2RY12 demonstrate (**Di**) elevated numbers of microglia that have (**Dii**) decreased expression of P2RY12, an indicator of increased reactive microglia, in EAE mice (data normalized to controls). (**E**) Immunostaining against glial fibrillary acidic protein (GFAP). There is a (**Ei**) significant increase in GFAP, indicative of reactive astrocytes in the LGN from EAE mice. (**D,E**) P2RY12 and GFAP stains further show pronounced changes in cell morphology of both cell types. (**B-E**) n=4. Scale bars, 20 μm. Data represent mean ± SEM, significant differences with $^*P < 0.05$, $^{**}P < 0.01$, $^{****}P < 0.0001$, t-test.

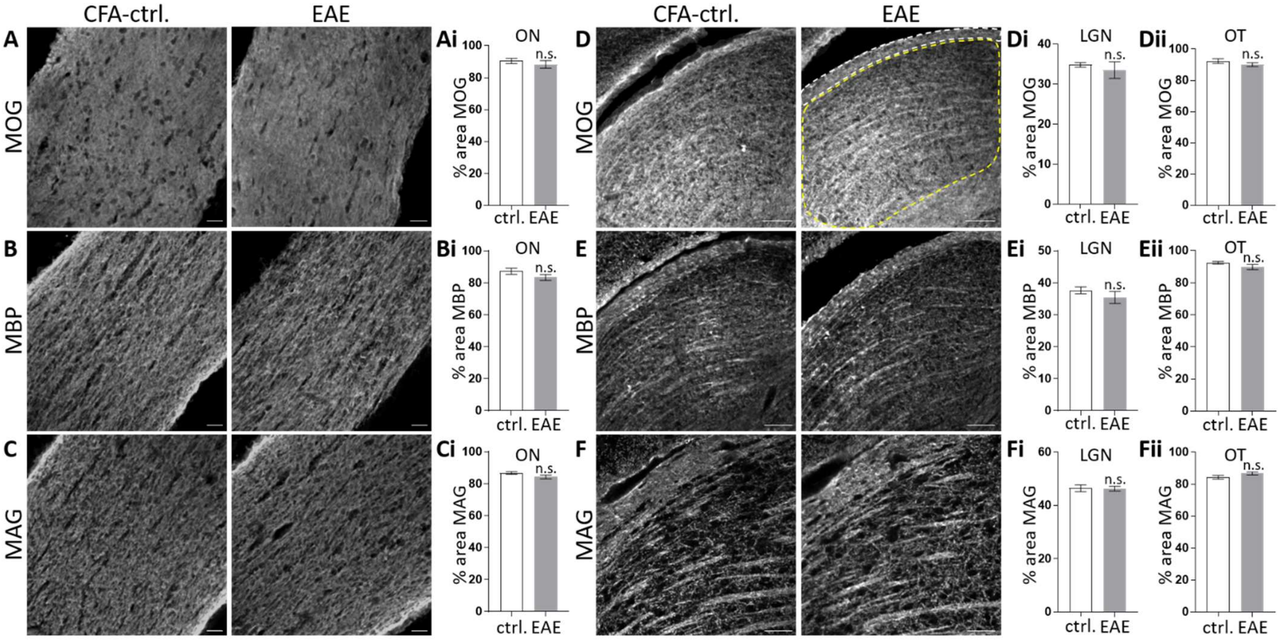


**Figure S3: No significant changes in myelin protein in the optic nerve, LGN and optic tract at early stages of EAE in the mouse.** (**A-F**) Analysis of myelin proteins from the same CFA-control and EAE mice as shown in Figs. 2-3 (clinical score=1.35 ±0.35). Representative images of the (**A-C**) optic nerve (ON), and the (**D-F**) LGN (outlined in yellow)/optic tract (OT, outlined in white) from CFA-control and EAE mice immunostained for (**A,D**) MOG, (**B,E**) MBP, and (**C,F**) MAG. There are (**Ai-Fii**) no significant changes in myelin protein in any region assessed within the retinogeniculate circuit at early stages of EAE. n=4. Scale bars, (**A-C**) 20 µm, (**D-F**) 100 µm. Data represent mean ± SEM, no significant differences (n.s.), t-test.

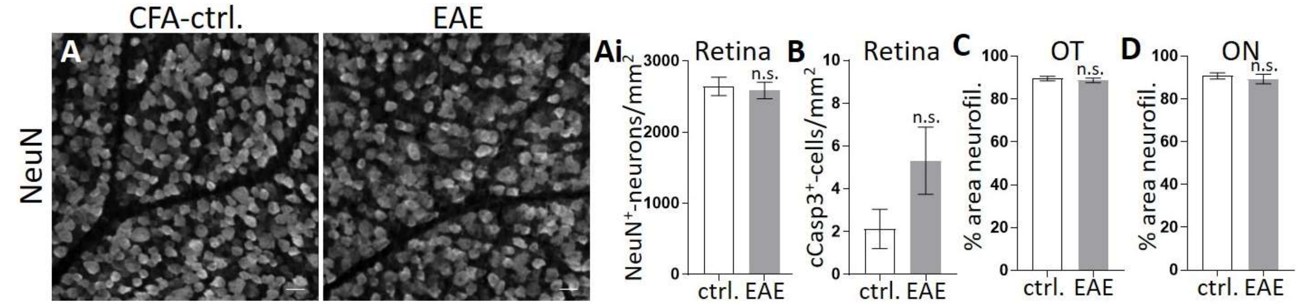


**Figure S4: No detectable neuron loss within the ganglion cell layer of the retina and no axonal loss in the optic tract and optic nerve at early stages of EAE.** (**A-D**) Analysis of the (**A,B**) retina, (**C**) optic tract (OT) and (**D**) optic nerve (ON) of the same CFA-control and EAE mice shown in Figs. 2-3 (clinical score=1.35 ±0.35). (**A**) Representative images of whole mount retina from CFA-control and EAE mice. Anti-NeuN staining was used to label neurons in the ganglion cell layer of the retina. There are no significant differences in the density of (**Ai**) $NeuN^+$-neurons and (**B**) cleaved caspase $3^+$-apoptotic cells between groups. (**C,D**) Further quantification of $neurofilament^+$-axons in the (**C**) OT and (**D**) ON reveals no alterations in the density of axons at early stages of EAE. (**A-D**) n=4. Scale bars, 20 µm. Data represent mean ± SEM, no significant differences (n.s.), t-test.

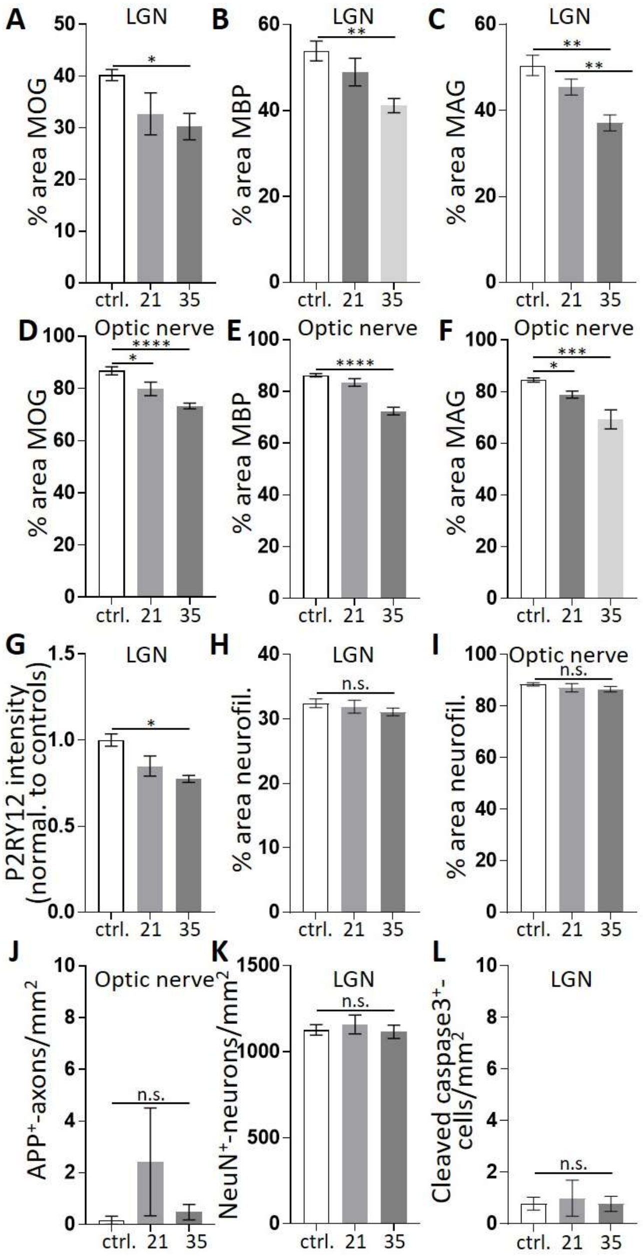


**Figure S5: No significant axon degeneration or neuron cell death at early and late phases of demyelination in the LGN of DTA mice.** (**A-L**) Further analysis of adjacent tissue sections from the same control and DTA mice shown in Fig. 4. (**A-F**) Indicative of demyelination, there is a significant decrease in myelin proteins (**A,D**) myelin oligodendrocyte glycoprotein (MOG), (**B,E**) myelin basic protein (MBP) and (**C,F**) myelin-associated glycoprotein (MAG) in the (**A-C**) LGN and (**D-F**) optic nerve in 35dpi DTA mice compared to controls (ctrl.). (**G**) Indicative of microgliosis, there is decreased intensity of anti-P2RY12 immunostaining in 35 dpi DTA mice (data normalized to controls). (**H-J**) There is no significant change in (**H,I**) neurofilament (neurofil.)-labeled axons in the (**H**) LGN or (**I**) optic nerve and no change in (**J**) APP levels in the optic nerve in DTA mice compared to controls. (**K,L**) There is also no significant change in the density of (**K**) $NeuN^+$- or (**L**) cleaved caspase $3^+$-cells in the LGN between groups. (**A-L**) n=9 ctrl./5 DTA-21dpi/6 DTA-35dpi. Data represent mean ± SEM, significant differences with $^*P < 0.05$, $^{**}P < 0.01$, $^{***}P < 0.001$, $^{****}P < 0.0001$, ANOVA with Tukey's *post hoc* test.

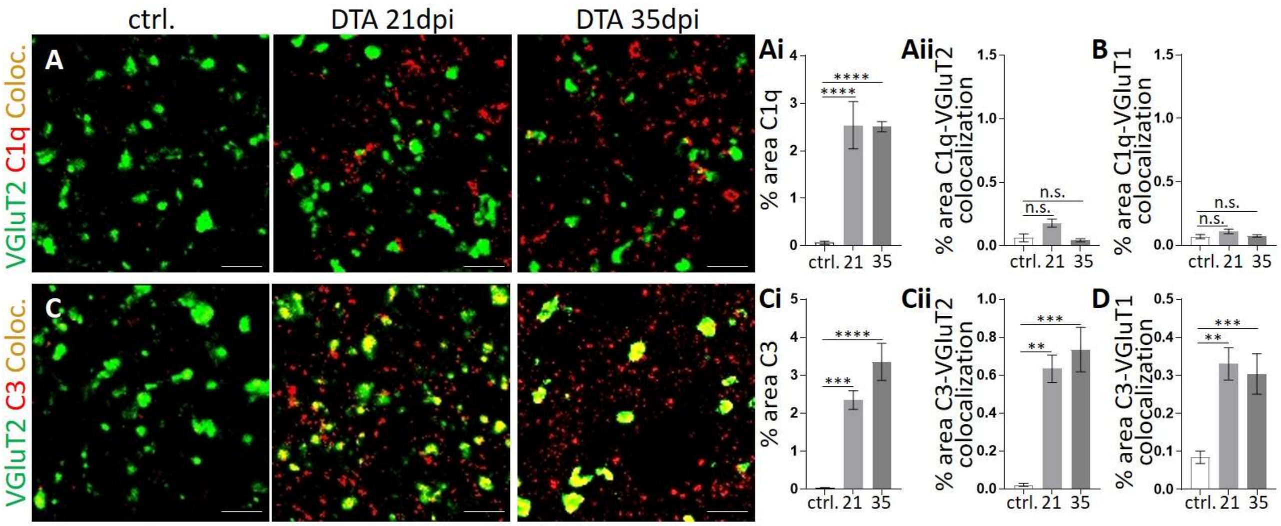


**Figure S6: Complement component C3, but not C1q, localizes to synapses in DTA mice.** (**A,C**) Representative confocal images from adjacent tissue sections of the LGN of control and DTA mice shown in Fig. 4 immunostained against retinogeniculate presynaptic terminal marker VGluT2 (green) and complement component (**A**) C1q or (**C**) C3 (red). There is a (**Ai**) significant increase in C1q in 21dpi and 35dpi DTA mice compared to controls. However, (**Aii,B)** C1q does not localize to presynaptic compartments. There is a (**Ci**) significant increase in C3 in the LGN of 21dpi and 35dpi DTA mice, which (**Cii,D**) colocalizes with presynaptic markers. (**A-D**) n= 5 ctrl./4 DTA. (**A,C**) Scale bars, 5 µm. Data represent mean ± SEM, significant differences with **P < 0.01, ***P < 0.001, ****P < 0.0001, ANOVA with Tukey's *post hoc* test.

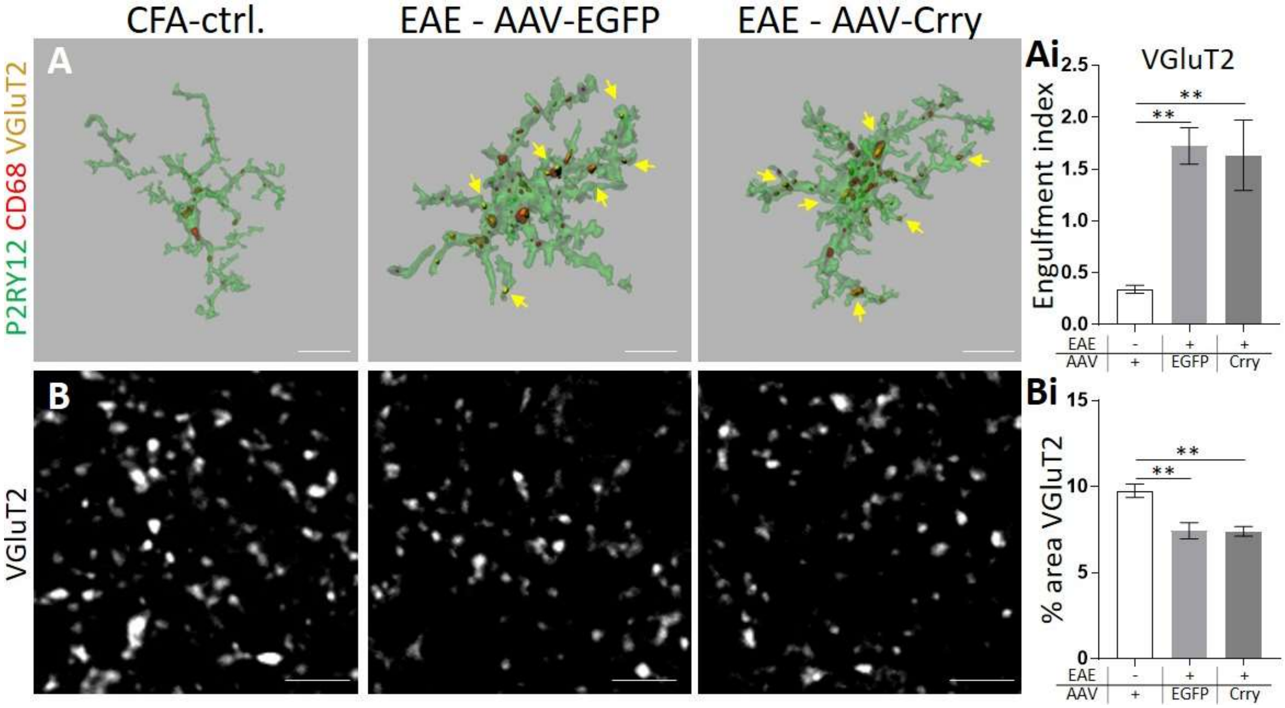


**Figure S7: Microglial engulfment and synapse loss in the somatosensory cortex following EAE is not blocked by retinal delivery of AAV-Crry.** (**A-B**) Analysis of layer IV of the somatosensory cortex of EAE and control treated mice shown in Fig. 6. (**A**) Representative 3D-surface rendering of P2RY12⁺-microglia, CD68-labeled lysosomes, and engulfed VGluT2⁺-retinogeniculate inputs within CD68-labeled lysosomes. Scale bars, 10 µm. (**Ai**) Microglia engulf significantly more VGluT2⁺-retinogeniculate presynaptic inputs (arrows) within CD68⁺-lysosomes in EAE mice injected with AAV-EGFP compared to CFA-controls (ctrl.), but this is not blocked by intravitreal delivery of AAV-Crry. (**B**) Representative confocal images of layer IV of the somatosensory cortex of the same animals immunostained against presynaptic terminal marker VGluT2. (**Bi**) There are significantly reduced densities of presynaptic VGluT2 in EAE mice injected with AAV-EGFP compared to CFA-controls (ctrl.), but this is not attenuated upon intravitreal delivery of AAV-Crry. Scale bars, 5 µm. (**A,B**) n=5. Data represent mean ± SEM, significant differences with $^{**}P < 0.01$, ANOVA with Tukey's *post hoc* test.

**Table S1: Human source information.**

| Patient No. | Sex | Age (years) | Diagnosis | MS Duration | Postmortem interval (hours) | Cause of Death |
|---|---|---|---|---|---|---|
| #1. | Male | 59 | primary progressive multiple sclerosis | 21 years | 9 | Embolic brain stem strokes |
| #2. | Female | 60 | progressive multiple sclerosis | 16 years | 7 | Sepsis (osteomyelitis, Clostridium difficile colitis, WBC>100.000) |
| #3. | Male | 76 | progressive multiple sclerosis | 34 years | 12 | Sepsis, chronic respiratory failure, aspiration pneumonia |
| #4. | Female | 78 | progressive multiple sclerosis | 54 years | 12 | Sepsis |
| #5. | Female | 78 | progressive multiple sclerosis | 24 years | 24 | Aspiration pneumonia |
| #6. | Female | 58 | control | N/A | 24 | Systemic sarcoidosis |
| #7. | Male | 66 | control | N/A | 33 | Hypertensive, atherosclerotic, and valvular cardiovascular disease |
| #8. | Female | 65 | control | N/A | 28 | Acute pulmonary thromboembolism due to deep vein thromboses of lower extremities following hypertensive and arteriosclerotic cardiovascular disease |
| #9. | Male | 84 | control | N/A | 28 | Multiple blunt impact injuries |
| #10. | Male | 64 | control | N/A | 50 | Hypertensive cardiovascular disease |

**Table S2: Marmoset source information.**

HHV6A/B= Human herpes virus 6 serotype A or B; SUP-T1 ctrl.= uninfected, virus-free human T-lymphoblast SUP-T1 cell line.

| Marmoset No. | Sex | Age at sacrifice (months) | Type | Experiment duration (weeks) | Visible lesion in the optic nerve/tract on *in vivo* MRI | Severity in the optic nerve/tract on *ex vivo* MRI | Laterality on *ex vivo* MRI |
|---|---|---|---|---|---|---|---|
| #1. Boozie | Male | 44 | EAE +HHV6B | 12 | Chiasm lesion visible | Severe | bilateral throughout, R > L |
| #2. Petra | Female | 38 | EAE | 18 | L nerve/chiasm visible | Severe | bilateral in chiasm only, L >>> R |
| #3. Stella | Female | 60 | EAE+SUP-T1 ctrl. | 23 | L&R Tract lesion visible | Mild | bilateral, L > R |
| #4. Pixie | Female | 38 | EAE | 15 | R chiasm lesion visible, L&R tract conspicuous | Medium | bilateral, L ≈ R |
| #5. Reggie | Male | 47 | EAE+SUP-T1 ctrl. | 134 | L&R Ant tract, R Post tract lesions visible | Medium | bilateral, L > R |
| #6. Michele | Female | 56 | EAE+SUP-T1 ctrl. | 19 | Conspicuous scattered lesions visible | Severe | bilateral throughout, L ≈ R |
| #7. Proton | Male | 39 | Naïve | N/A | N/A | None | N/A |
| #8. Laura | Female | 65 | Naïve+HHV6B | N/A | N/A | None | N/A |
| #9. Tom | Male | 80 | EAE | 61 | N/A | None | N/A |
| #10. Katie | Female | 53 | EAE+ HHV6A | 8 | N/A | None | N/A |
| #11. Beijing | Female | 31 | EAE+ HHV6B | 4 | N/A | Minimal | bilateral, L > R |
| #12. Edith | Female | 49 | EAE+ HHV6A | 7 | N/A | None | N/A |
